# Learning dynamical information from static protein and sequencing data

**DOI:** 10.1101/401067

**Authors:** Philip Pearce, Francis G. Woodhouse, Aden Forrow, Ashley Kelly, Halim Kusumaatmaja, Jörn Dunkel

## Abstract

Many complex processes, from protein folding and virus evolution to brain activity and neuronal network dynamics, can be described as stochastic exploration of a high-dimensional energy landscape. While efficient algorithms for cluster detection and data completion in high-dimensional spaces have been developed and applied over the last two decades, considerably less is known about the reliable inference of state transition dynamics in such settings. Here, we introduce a flexible and robust numerical framework to infer Markovian transition networks directly from time-independent data sampled from stationary equilibrium distributions. Our approach combines Gaussian mixture approximations and self-consistent dimensionality reduction with minimal-energy path estimation and multi-dimensional transition-state theory. We demonstrate the practical potential of the inference scheme by reconstructing the network dynamics for several protein folding transitions, gene regulatory network motifs and HIV evolution pathways. The predicted network topologies and relative transition time scales agree well with direct estimates from time-dependent molecular dynamics data, stochastic simulations and phylogenetic trees, respectively. The underlying numerical protocol thus allows the recovery of relevant dynamical information from instantaneous ensemble measurements, effectively alleviating the need for time-dependent data in many situations. Owing to its generic structure, the framework introduced here will be applicable to high-throughput RNA and protein sequencing datasets and future cryo-electron-microscopy data, and can guide the design of new experimental approaches towards studying complex multiphase phenomena.

## INTRODUCTION

Energy landscapes encapsulate the effective dynamics of a wide variety of physical, biological and chemical systems^1–3^. Well-known examples include a myriad of biophysical processes^4–11^, multiphase systems^2^, thermally activated hopping in optical traps^12,13^, chemical reactions^1,14^, brain neuronal expression^15^, cellular development^16–20^ and social networks^21,22^. Energetic concepts have also been connected to machine learning^23^ and to viral fitness landscapes, where pathways with the lowest energy barriers may explain typical mutational evolutionary trajectories of viruses between fitness peaks^24,25^. Recent advances in experimental techniques including cryo-electron microscopy (cryo-EM)^4,26–30^ and single-cell RNA sequencing^31^, as well as new online social interaction datasets^32,33^, are producing an unprecedented wealth of high-dimensional instantaneous snapshots of biophysical and social systems. Although much progress has been made in dimensionality reduction^34–37^ and the reconstruction of effective energy landscapes in these settings^4,17,20,22,30^, the problem of inferring dynamical information such as protein folding or mutation pathways and rates from instantaneous ensemble data remains a major challenge.

To address this practically important question, we introduce here an integrated computational framework for identifying metastable states on reconstructed high-dimensional energy landscapes and for predicting the relative mean first passage times (MFPTs) between those states, without requiring explicitly time-dependent data. Our inference scheme employs an analytic representation of the data based on a Gaussian mixture model (GMM)^38^ to enable efficient identification of minimum-energy transition pathways^39–41^. We show how the estimation of transition networks can be optimized by reducing the dimension of a high-dimensional landscape while preserving its topology. Our algorithm utilizes experimentally validated analytical results^12,13^ for transition rates^1,42–44^. Thus, it is applicable whenever the time-evolution of the underlying system can be approximated by a Fokker-Planck-type Markovian dynamics, as is the case for a wide range of physical, chemical and biological processes^1,43,45^.

Specifically, we illustrate the practical potential by inferring protein folding transitions, state-switching in gene regulatory networks and HIV evolution pathways. Current standard methods for coarse-graining the conformational dynamics of biophysical structures^46–48^ typically estimate Markovian transition rates from time-dependent trajectory data in large-scale molecular dynamics simulations^49,50^. By contrast, we show here that protein folding pathways and rates can be recovered without explicit knowledge of the time-dependent trajectories, provided the system is sufficiently ergodic and equilibrium distributions are sampled accurately. Furthermore, we show that the dynamics of state-switching or phenotype-switching in gene regulatory networks^51^ can be inferred directly from static snapshots of protein abundances in regimes where deterministic modeling only captures a single steady state^52,53^. The agreement of our inferred results with two separate sets of time-dependent measurements suggests that the inference of complex transition networks via reconstructed energy landscapes can provide a viable and often more efficient alternative to traditional time series estimates, particularly as new experimental techniques will offer unprecedented access to high-dimensional ensemble data.

## RESULTS

### Minimum-energy-path (MEP) network reconstruction

The equilibrium distribution *p*(x) of a particle diffusing over a potential energy landscape *E*(x) is the Boltzmann distribution *p*(x) = exp [−*E*(x)/*k*_*B*_*T*]/*Z*, where *k*_*B*_ is the Boltzmann constant, *T* is the temperature and *Z* is a normalization constant. Given the probability density function (PDF) *p*(x), the effective energy can be inferred from

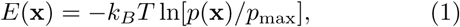

where *p*_max_ is the maximum value of the PDF, included to fix the minimum energy at zero. Our goal is to estimate the MFPTs between minima on the landscape using only sampled data. We divide this task into three steps, as illustrated in Fig. 1 for test data (Supplementary Information). In the first step, we approximate the empirical PDF by using the expectation maximization algorithm to fit a Gaussian mixture model (GMM) in a space of sufficiently large dimension *d* (Methods, Fig 1A). Mixtures with a bounded number of components can be recovered in time polynomial in both *d* and the required accuracy^54^. The resulting GMM yields an analytical expression for *E*(x) via Eq. (1).

**FIG. 1.**
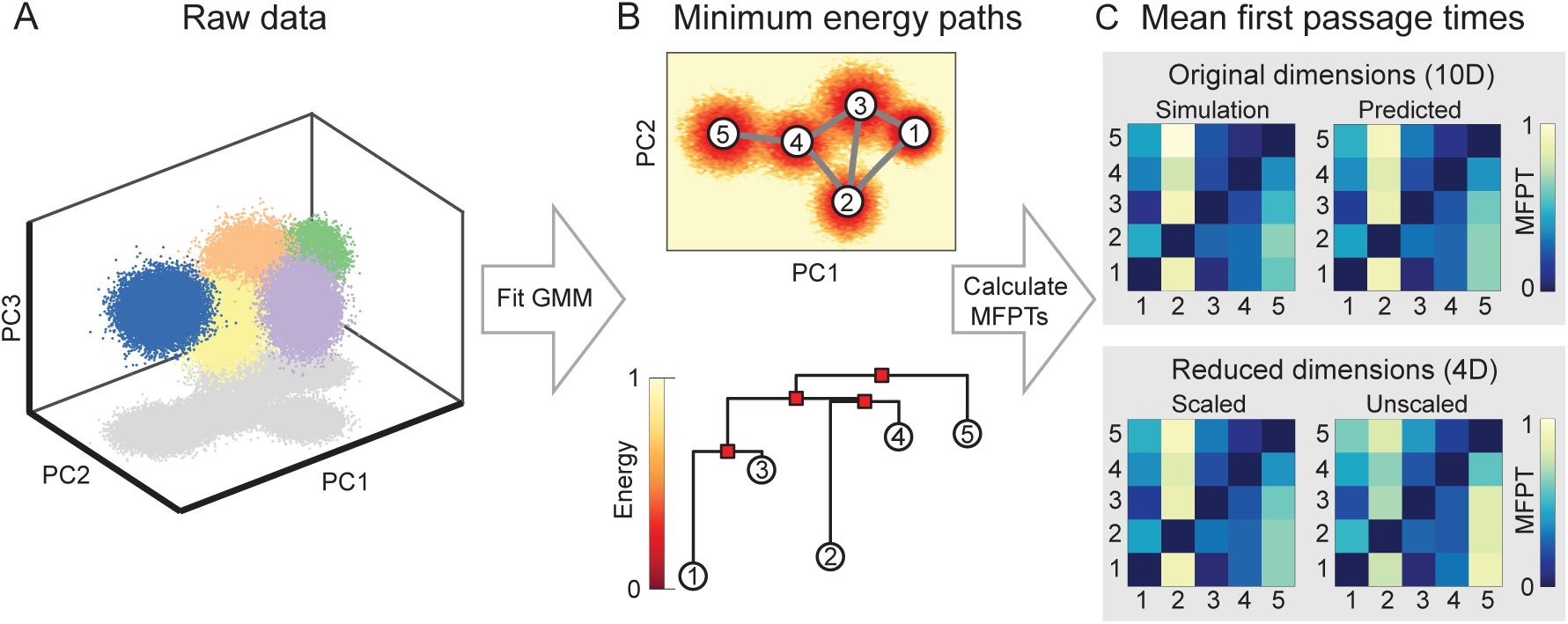
Inference scheme for estimating transition networks and mean first passage times (MFPTs) from a stationary sample set, demonstrated on test data generated from a Gaussian Mixture Model (GMM; Supplementary Information). *(A)* Inputs are the instantaneously measured data, sampled here from a 10-dimensional GMM with 5 Gaussians, plotted in the first 3 principal components (PCs). *(B)* Top: A GMM is fit to the samples to construct the empirical distribution which is then converted to the energy landscape using Eq. (1). Background color indicates the projection of the empirical energy landscape onto the first two PCs. Minimum energy paths (MEPs, grey lines) between minima 1–5 on the landscape are calculated using the NEB algorithm (Supplementary Information). Bottom: Disconnectivity graph illustrating minima on the energy landscape (circles) and saddle points between them (squares). *(C)* A Markov state model (MSM) is constructed with transition rates given by Eq. (2) and solved to predict the MFPTs between discrete states (top right; Methods). MFPTs predicted by the MSM agree with direct estimates from Brownian dynamics simulations in the inferred energy landscape (top left; Supplementary Information). MFPTs calculated in a reduced 4-dimensional space using the scaling given in Eq. (3) recover the MFPTs accurately (bottom left). Without the appropriate scaling, the predicted MFPTs are inaccurate (bottom right).

In the second step, the inferred energy landscape *E*(x) is reduced to an MEP network whose nodes (states) are the minima of *E*(x) (Fig. 1B top). Each edge represents an MEP that connects two adjacent minima and passes through an intermediate saddle point (Fig. 1B). The MEPs are found using the nudged elastic band (NEB) algorithm^39,40^, which discretizes paths with a series of bead-spring segments (Supplementary Information).

### Markov state model (MSM)

Given the MEP network, the final step is to infer the rates for transitioning from a minimum *α* to an adjacent minimum *β*. Assuming overdamped Brownian dynamics, the directed transition *α → β* can be characterized by the generalized Kramers transition rate^1^

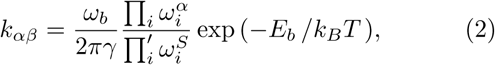

where *γ* is the effective friction, *E*_*b*_ is the energy difference between the saddle point *S* on the MEP (over the energy barrier) and the minimum *α*, 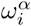 are the stable angular frequencies at the minimum *α*, while 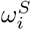 and *ω*_*b*_ are the stable and unstable angular frequencies at the saddle. Eq. (2) assumes isotropic friction but can be generalized to a tensorial form^1^ if anisotropies are relevant. In most practical applications, the error from assuming *γ* to be isotropic is likely negligible compared to other experimental noise sources. In principle, Eq. (2) can be refined further by including quartic (or higher) corrections to the prefactor *ω*_*b*_/*γ* to account for details of the saddle shape^1^. Such corrections can be significant for GMMs (Supplementary Information).

Each edge (*αβ*) has two weights, *k_αβ_* and *k_βα_*, assigned to it. The rate matrix (*k_αβ_*) completely specifies the MSM on the network. Solving the MSM yields the matrix of pairwise mean first passage times (MFPTs) between states (Fig. 1C, Methods). In a simple two-state system, the MFPTs are determined up to a time scale by detailed balance, but for three or more states the influence of landscape topography and the associated state network topology (Methods) can lead to interesting hierarchical ordering of passage times. Identifying these hierarchies, and ways to manipulate them, is key to controlling protein folding or viral evolution pathways.

### Topology-preserving dimensionality reduction

To ensure that the inference protocol can be efficiently applied to larger systems with a high-dimensional energy landscape, we derive a general method for reducing the dimension *D* of an energy landscape while preserving its topology. A probability density function with *C* well-separated Gaussians in *D* dimensions can be projected onto the *d* = *C* – 1 dimensional hyperplane spanning the Gaussian means using principal component analysis (PCA). In practice, it suffices to choose *C* to be larger than the number of energy minima if their number is not known in advance. Reduction to fewer than *d* = *C* – 1 dimensions in general does not allow a correct recovery of the MFPTs.

To preserve the topology under such a transformation – which is essential for the correct preservation of energy barriers and MEPs in the reduced-dimensional space – one needs to rescale GMM components in the low-dimensional space depending on the covariances of the Gaussians in the *D* – *d* neglected dimensions (Fig. 1C). Explicitly, one finds that within the subspace spanned by the retained principal components (Supplementary Information)

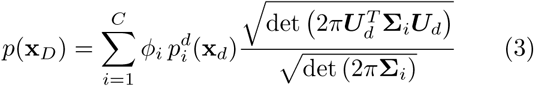

as long as *p* satisfies certain minimally-restrictive conditions (Supplementary Information). Here, ***U***_*d*_ denotes the first *d* = *C* – 1 columns of the matrix of sorted eigenvectors ***U*** of the covariance matrix of the Gaussian means, and *ϕ*_*i*_, 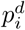 and **Σ**_*i*_ are the mixing components, reduced-dimensional PDF and the covariance matrix of each individual Gaussian in the mixture, respectively (Supplementary Information). Neglecting the determinant scale-factors in Eq. (3), as is often done when GMM models are fitted to PCA-projected data, leads to inaccurate MFPT estimates (Fig. 1C, bottom). Note that Eq. (3) does not represent inversion of the transformation performed on the data by PCA, unless all *D* dimensions are retained; if some dimensions are neglected, Eq. (3) represents a rescaling of the marginal distribution in the retained dimensions to reconstruct the probability density function in the original dimension. In other words, the transition rates are best recovered from the conditional – not marginal – distributions, which are given by Eq. (3) up to a constant factor that does not affect energy differences.

Dimensionality reduction can substantially improve the efficiency of the NEB algorithm step: when the MEPs in the reduced *d*-dimensional space have been computed, the identified minima and saddles can be transformed back into the original data dimension *D* to calculate the Hessian matrices at these points, allowing Kramers’ rates to be calculated as usual (Fig. 1C, Supplementary Information). Alternatively, in specific situations where the MEPs lie outside the hyperplane spanning the means (Supplementary Information), the MEP in the reduced *d*-dimensional space can be transformed back to the *D*-dimensional space and used as an initial condition in that space, significantly reducing computational cost. These results present a step towards a general protocol for identifying reaction coordinates or collective variables for projection of a high-dimensional landscape onto a reduced space while quantitatively preserving the topology of the landscape.

### Protein folding

To illustrate the vast practical potential of the above scheme, we demonstrate the successful recovery of several protein folding pathways, using data from previous large-scale molecular dynamics (MD) simulations^49^. The protein trajectories, consisting of the time-dependent coordinates of the alpha carbon backbone, were pre-processed, subsampled by a factor of 5, treated as a set of static equilibrium measurements, and reduced in dimension before fitting a GMM (Methods). As is typical for high-dimensional parameter estimation with few structural assumptions, the fitting error due to a finite sample size *n* in *d* dimensions scales approximately as 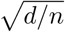 (Supplementary Information); see Refs. 55–57 for advanced techniques tackling sample size limitations. Here, *d <* 10 so the sample size *n* ∼ 10^5^ suffices for effective recovery; indeed, our results were found to be robust for trajectories further subsampled by up to a factor of 25, leaving around 500 samples per Gaussian (Fig. S3).

For each of the four analyzed proteins Villin, BBA, NTL9 and WW, the reconstructed energy landscapes reveal multiple states including a clear global minimum corresponding to the folded state (Fig. 2A,B). To estimate MFPTs, we determined the effective friction *γ* in Eq. (2) for each protein from the condition that the line of best fit through the predicted vs. measured MFPTs has unit gradient. Although not usually known, *γ* could in principle be calculated by comparing MD simulations with experimental data. Our MFPT predictions agree well with direct estimates (Supplementary Information) from the time-dependent MD trajectories (Fig 2C). Detailed analysis confirms that the MFPT estimates are robust under variations of the number of Gaussians used in the mixture (Fig. S1). Also, the estimated MEPs are in good agreement with the typical transition paths observed in the MD trajectories (Fig. S2).

**FIG. 2.**
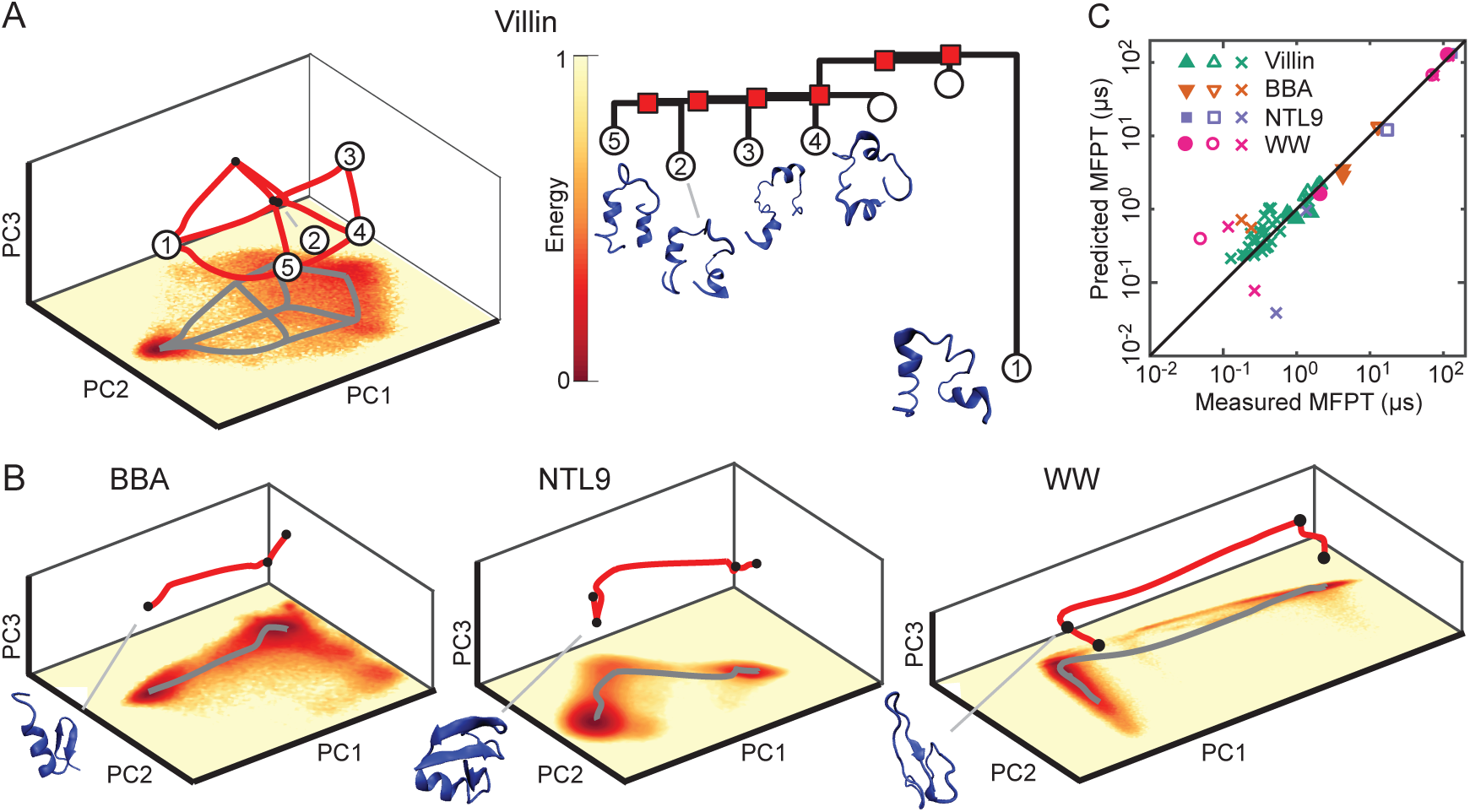
Reconstructed MEP networks for protein folding transitions, and comparison of predicted MFPTs with direct estimates from molecular dynamics (MD) simulations (Supplementary Information). *(A)* Left: Low energy states and transition network in the first three principal components (PCs) for Villin including predicted transition paths between states (red lines); bottom coloring shows two-dimensional projection of the empirical energy landscape onto the first two PCs. Right: Associated disconnectivity graph and illustrations of the five lowest energy states, with state 1 corresponding to the folded state. *(B)* Low energy states, transition paths and empirical energy landscape for BBA, WW and NTL9 proteins, and sketches of their folded states. *(C)* Predicted MFPTs agree well with estimates from MD simulations when energy minima are well separated and become less accurate for fast transitions with small MFPTs. Filled shapes correspond to transitions ending in the unfolded state and unfilled shapes correspond to transitions ending in the folded state (Supplementary Information), for Villin, BBA, NTL9 and WW. Crosses correspond to transitions between intermediate states.

### Gene regulatory networks

Next, we demonstrate the ability of our protocol to infer state-switching pathways in multistable gene regulatory networks. Using a Gillespie stochastic simulation algorithm (SSA; Methods), we simulated three repressilator-type gene regulatory network motifs^58^ with self-activation. Gene network motifs with features such as these have been studied extensively in recent years owing to their ability to exhibit precise oscillations^59^ and to their possible importance in the determination of multiple cell fates^60^ in the appropriate parameter regimes, although the role of noise in such networks is not well understood. In our simulated gene networks, each gene encodes a protein that activates the expression of its associated gene and represses another, with *D* = 2, 3 and 4 dimensions at low molecule numbers (Fig. 3A, Supplementary Information). The energy landscapes reconstructed from the simulation datasets in protein molecule-number space (with time-dependence removed) revealed multiple metastable states for each network (Fig. 3B). Broadly, we found each state to correspond to a mixture of low and high abundances for each separate protein, with the two most common states in *D* = 4 dimensions consisting of two abundant and two depleted proteins (Fig. 3B). In agreement with previous studies^52,53^, the identified metastable states were not recovered from deterministic simulations of the governing ordinary differential equations (Supplementary Information), but could only be identified directly from the stochastic data (Fig. 3A,B). We determined the effective friction *γ* in Eq. (2) for each *D* as in the protein example. The predicted MFPTs between each metastable state were found to be accurate in comparison to time-dependent measurements (Fig. 3C), demonstrating the utility of our protocol for gene regulatory network datasets and, more generally, energy landscapes in discrete spaces.

**FIG. 3.**
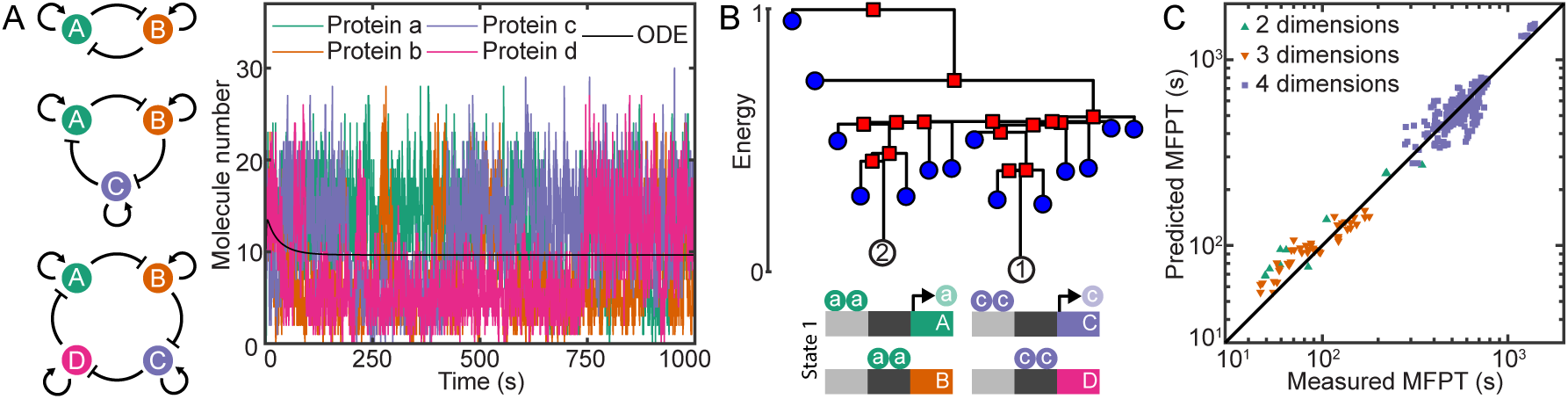
Reconstructed MEP networks for multistable gene regulatory networks, and comparison of predicted MFPTs with direct estimates from stochastic simulations (Methods). *(A)* We performed simulations of three repressilator-type gene regulatory network motifs with self-activation (left), consisting of two (top), three (middle) and four (bottom) genes, denoted A, B, C, and D. In the stochastic simulations (right), the numbers of each protein fluctuate between metastable states, but deterministic simulations of the system of ordinary differential equations (ODEs) at low molecule numbers are not able to identify the states, instead converging to a single steady state where all protein numbers are equal. *(B)* Disconnectivity graph and illustration of the identified lowest energy state for the four-dimensional system. In the lowest energy state (State 1), larger numbers of proteins a and c are present, leading to the activation of their associated genes and repressing the expression of proteins b and d. The next lowest energy state (State 2) is the converse scenario, with large numbers of proteins b and d present. *(C)* Predicted MFPTs agree well with those calculated from the time-dependent stochastic simulations for all three of the network motifs.

### Viral evolution

As a final proof-of-concept application, we demonstrate that our inference scheme recovers the expected evolution pathways between HIV sequences as well as the key features of a distance-based phylogenetic tree (Fig. 4). To this end, we reconstructed an effective energy landscape from publicly available HIV sequences sampled longitudinally at several points in time from multiple patients^61^, assuming that the frequency of an observed genotype is proportional to its probability of fixation and that the high-dimensional discrete sequence space can be projected onto a continuous reduced-dimensional phenotype space (Fig. 4A; Supplementary Information). First, a Gaussian was fit to each patient and then combined in a GMM with equal weights, to avoid bias in the fitness landscape towards sequences infecting any specific patient (Supplementary Information). Thereafter, we applied our inference protocol to reconstruct the effective energy landscape, transition network (Fig. 4B) and disconnectivity graph (Fig. 4C), where each state is associated to a separate patient. As expected, states corresponding to patients infected with different HIV subtypes are not connected by MEPs (Fig 4A,B). The disconnectivity graph reproduces the key features of a coarse-grained patient-level representation of the phylogenetic tree (Fig. 4C). Using our inference scheme, vertical evolution in the tree can be tracked along the minimum energy paths in a reduced-dimensional sequence space (Fig. 4B). The energy barriers, represented by the lengths of the vertical lines in the disconnectivity graph (Fig. 4C), provide an estimate for the relative likelihood of evolution to fixation via point mutations between fitness peaks (energy minima). If mutation rates are known, the MEPs can also be used to estimate the time for evolution to fixation from one fitness peak to another^62^.

**FIG. 4.**
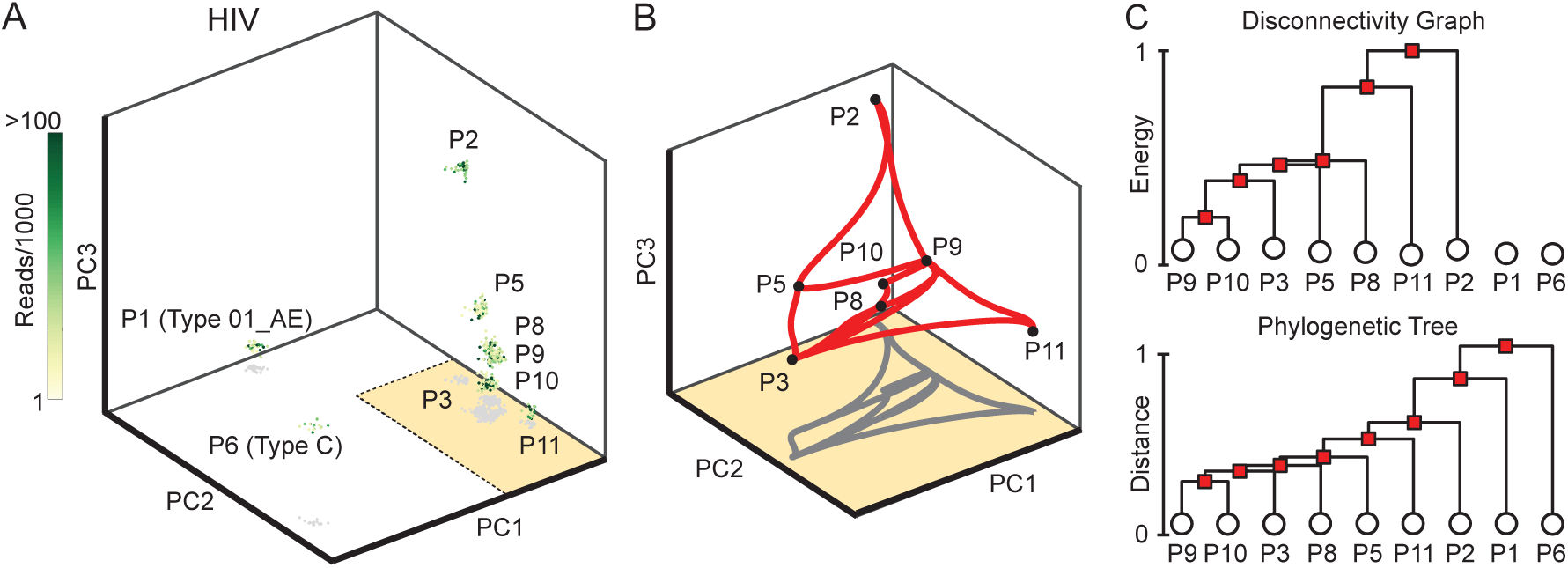
Minimum energy paths (MEPs) on viral fitness landscapes reconstructed from publicly available HIV sequencing data^61^. *(A)* Longitudinal samples of the HIV virus are binarized after multiple sequence alignment (Supplementary Information) and plotted in the first 3 PCs. Samples of the same HIV subtype are closer in PC-space. Patient labels correspond to those used in^61^. *(B)* MEPs between minima corresponding to patients infected with Type B HIV, plotted in the first 3 PCs. Paths between minima indicate likely evolutionary pathways. Minima corresponding to patients with Type 01 AE and Type C HIV were unconnected to the other minima. *(C)* Disconnectivity graph for connected minima, where vertical evolution frequency is assumed to be proportional to the normalized energy barriers (top). The disconnectivity graph reproduces the majority of the structure of a distance-based phylogenetic tree (bottom), where the lengths of vertical lines are proportional to the Jukes-Cantor sequence distance (scaled to [0, 1]).

## DISCUSSION

### Preserving landscape topology under dimensionality reduction

Finding the appropriate number of collective macro-variables to describe an energy landscape is a generic problem relevant to many fields. For example, although some proteins can be described through effective one-dimensional reaction coordinates^6,9,63,64^, the accurate description of their diffusive dynamics over the full microscopic energy landscape requires many degrees of freedom^65,66^. Whenever dynamics are inherently high-dimensional, topology-preserving dimensionality reduction can enable a much faster search of the energy landscape for minima and MEPs. In practice, data dimension is often reduced with PCA or similar methods before constructing an energy landscape^66–73^. The extent to which commonly used dimensionality reduction techniques alter MEP network topology or quantitatively preserve energy barriers is not well understood. Eq. (3) suggests that reducing dimensions using PCA should not introduce significant errors if the variance of the landscape around each state (energy minimum) in the neglected dimensions is similar. For instance, we found that the protein folding data could be reduced to five dimensions while maintaining accuracy (Fig. S1), although additional higher energy states may become evident in higher dimensions. As an alternative to using Eq. (2) in the last stage of our approach, a method such as maximum caliber^74–76^, which does not take the derivatives of landscape topology into account, could be supplied with the sizes of the energy barriers and used to infer MFPTs. However, we found that owing to the dependence of the MFPTs on the prefactors in Eq. (2) for different transitions, this technique could not recover all transition rates accurately for either proteins or gene regulatory networks (Fig. S4). Overall, our theoretical results demonstrate the benefits of combining an analytical PDF with a linear dimensionality reduction technique so that the neglected dimensions can be accounted for explicitly.

### Biological and biophysical applications

Rapidly advancing imaging techniques, such as cryogenic electron microscopy (cryo-EM), will allow many snapshots of biophysical structures to be taken at the atomic level in the near future^4,26–30^. A biologically and biophysically important task will be to infer dynamical information from such instantaneous static ensemble measurements. The protein folding example in Fig. 2 suggests that the framework introduced here can help overcome this major challenge; in principle the framework requires only the pairwise distances between recognizable features of the protein as input (here we used the carbon alpha coordinates). Another promising area of future application is the analysis of single-cell RNA-sequencing data quantifying the expression within individual cells^31^. Related to this application, Fig. 3 demonstrates that our protocol recovers state-switching pathways in multistable gene regulatory networks, which are thought to underlie cell fate decisions. These results are most relevant in low-molecule-number regimes, in which noise is known to be an important factor^77^. In relevant recent work, an effective non-parametric energy landscape of single-cell expression snapshots was inferred using the Laplacian of a k-nearest neighbor graph on the data, allowing lineage information to be derived via a Markov chain^19^. The GMM-based framework here provides a complementary parametric approach for reconstructing faithful low-dimensional transition state dynamics from such high-dimensional data.

Furthermore, the proof-of-concept results in Fig. 4 suggests that our inference scheme for Markovian network dynamics can be useful for studying viral and bacterial evolution, which are often modeled as movements through a series of DNA or protein sequences^78^. The fitness landscape of an organism in sequence space is analogous to the negative of an effective energy landscape. The process of fixation by a succession of mutants in a population, whereby each mutant replaces the previous lineage as the population’s most recent common ancestor, has been modeled as a Markov process^79^. Successive sweeps to fixation have been observed in long-term evolution experiments, promising groundbreaking data for future analysis as whole-genome sequencing technologies improve^80^.

### Outlook and extensions

The inference protocol opens the possibility to analyze previously intractable multi-phase systems: many high-dimensional physical, chemical and other stochastic processes can be described by a Fokker-Planck dynamics^1^, with phase equilibria corresponding to maxima of the stationary distribution. By taking near-simultaneous measurements of many subsystems within a large multistable Fokker-Planck system, the above scheme allows the inference of coexisting equilibria and transition rates between them. Other possible applications may include neuronal expression^15^ and social networks^21,22,32,33^, which have been described in terms of effective energy landscapes.

While we focused here on normal white-noise diffusive behavior, as is typical of protein folding dynamics, the above ideas can in principle be generalized to other classes of stochastic exploration processes. Such extensions will require replacing Eq. (2) through suitable generalized rate formulas, as have been derived for correlated noise^1,81^. Conversely, the present framework provides a means to test for diffusive dynamics: if the MFPTs of an observed system differ markedly from those inferred by the above protocol, then either important degrees of freedom have not been measured; the system is out of equilibrium on measurement time scales; or the system does not have Brownian transition statistics, necessitating further careful investigation of its time dependence.

To conclude, the conformational dynamics of biophysical structures such as viruses and proteins, and the state-switching dynamics of noisy gene regulatory networks, are characterized by their metastable states and associated transition networks, and can often be captured through Markovian models. Current experimental techniques, such as cryo-EM or RNA-sequencing, provide limited dynamical information. In these cases, transition networks must be inferred from structural snapshots. Here, we have introduced and tested a numerical framework for inferring Markovian state-transition networks via reconstructed energy landscapes from high-dimensional static data. The successful application to protein folding, gene regulatory network and viral evolution pathways illustrates that high-dimensional energy landscapes can be reduced in dimension without losing relevant topological information. Generally, the inference scheme presented here is applicable whenever the dynamics of a high-dimensional physical, biological or social system can be approximated by diffusion in an effective energy landscape.

## METHODS

### Population landscapes

A Gaussian mixture model (GMM) was used to represent the probability density function (PDF), or population landscape, of samples. The PDF at position x of a GMM with *C* mixture components in *d* dimensions is

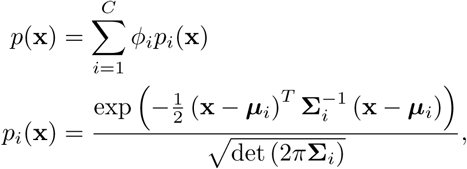

where *ϕ*_*i*_ are the weights of each component, ***µ***_*i*_ are the means and **Σ**_*i*_ are the covariance matrices. More details on GMMs and how they were fit to data is given in the Supplementary Information.

### Mean first passage times

We form a discrete-state continuous-time Markov chain on states given by the minima of the energy landscape. For a pair of states *α* and *β* directly connected by a minimum-energy pathway via a saddle, we approximate the transition rate *α → β* by the Kramers rate *k*_*αβ*_ in Eq. (2), while if *α* and *β* are not directly connected we set *k*_*αβ*_ = 0. Given these rates, the Markov chain has generator matrix *M*_*αβ*_ where *M*_*αβ*_ = *k*_*αβ*_ for *α* ≠ *β* and *M*_*αα*_ = −∑_*β*:*β*≠*α*_ *k*_*αβ*_. Then the matrix *τ*_*αβ*_ of MFPTs (hitting times) for transitions *α → β* satisfies

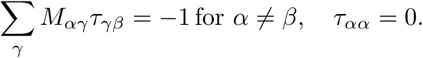

### Protein data pre-processing

Protein folding trajectories were obtained from all-atom molecular dynamics (MD) simulations performed by D.E. Shaw Research^49^. Data was subsampled by a factor of 5 to reduce the size. For some proteins, residues at the flexible tails of proteins were removed from the dataset to reduce noise. Pairwise distances between carbon alpha atoms on the protein backbone were taken, with a cut off of 6-8 Å, depending on the size of the protein; any distance above the threshold was taken to be equal to the threshold. This vector of pairwise distances was used as input to principal component analysis (PCA) to reduce dimension. The first five principle components of the protein data were found to be sufficient for inference of energy landscapes and transition networks (Fig. S1).

### Gene regulatory network simulations

Gene regulatory network motifs were simulated using a Gillespie stochastic simulation algorithm (SSA) in the SimBiology toolbox in Matlab. A full list of the reactions simulated for each motif, as well as the values of the parameters used, is given in the Supplementary Information and in the simulation code, which is available from Github (https://github.com/philip-pearce/learning-dynamical).

### Code availability

The source code used in this study to learn a dynamical transition network and mean first passage times from a Gaussian mixture model is publicly available from Github (https://github.com/philip-pearce/learning-dynamical). Also included are all data processing codes required to convert the raw data used in this study into the appropriate format.

### Data availability

Two publicly available datasets were used in this study. Protein folding trajectories^49^ are available from D.E. Shaw Research (https://www.deshawresearch.com/). HIV sequences^61^ are available from https://hiv.biozentrum.unibas.ch/.

## ACKNOWLEDGMENTS

We thank D.E. Shaw Research for protein folding trajectories and Stefano Piana-Agostinetti of D.E. Shaw Research for helpful discussions. This work was supported by the Royal Society International Exchanges award IE160909 (H.K. and J.D.) and Complex Systems Scholar Award from the James S. McDonnell Foundation (J.D.).

## References

1 Hänggi, P., Talkner, P. & Borkovec, M. Reaction-rate theory: Fifty years after Kramers. Rev. Mod. Phys. 62, 251–341 (1990).

2 Yukalov, V. Phase transitions and heterophase fluctuations. Phys. Rep. 208, 395–489 (1991).

3 Helbing, D. Traffic and related self-driven many-particle systems. Rev. Mod. Phys. 73, 1067–1141 (2001).

4 Dashti, A. et al. Trajectories of the ribosome as a Brownian nanomachine. Proc. Natl. Acad. Sci. USA 111, 17492–17497 (2014).

5 Chung, H. S., Piana-Agostinetti, S., Shaw, D. E. & Eaton, W. A. Structural origin of slow diffusion in protein folding. Science 349, 1504–1510 (2015).

6 Neupane, K., Manuel, A. P. & Woodside, M. T. Protein folding trajectories can be described quantitatively by one-dimensional diffusion over measured energy landscapes. Nat. Phys. 12, 700–703 (2016).

7 Hosseinizadeh, A. et al. Conformational landscape of a virus by single-particle X-ray scattering. Nat. Methods 14, 877–881 (2017).

8 Johnson, R. R., Kohlmeyer, A., Johnson, a. T. C. & Klein, M. L. Free energy landscape of a DNA-carbon nanotube hybrid using replica exchange molecular dynamics. Nano. Lett. 9, 537–541 (2009).

9 Best, R. B. & Hummer, G. Diffusive model of protein folding dynamics with Kramers turnover in rate. Phys. Rev. Lett. 96, 228104 (2006).

10 Zhou, L., Marras, A. E., Su, H. J. & Castro, C. E. Direct design of an energy landscape with bistable DNA origami mechanisms. Nano. Lett. 15, 1815–1821 (2015).

11 Fern, J., Lu, J. & Schulman, R. The energy landscape for the self-assembly of a two-dimensional DNA origami complex. ACS Nano 10, 1836–1844 (2016).

12 McCann, L. I., Dykman, M. & Golding, B. Thermally activated transitions in a bistable three-dimensional optical trap. Nature 402, 785–787 (1999).

13 Rondin, L. et al. Direct measurement of Kramers turnover with a levitated nanoparticle. Nat. Nanotechnol. 12, 1130–1133 (2017).

14 García-Müller, P. L., Borondo, F., Hernandez, R. & Benito, R. M. Solvent-induced acceleration of the rate of activation of a molecular reaction. Phys. Rev. Lett. 101, 178302 (2008).

15 Ezaki, T., Watanabe, T., Ohzeki, M. & Masuda, N. Energy landscape analysis of neuroimaging data. Phil. Trans. R. Soc. A 375, 20160287 (2016).

16 Corson, F. & Siggia, E. D. Geometry, epistasis, and developmental patterning. Proc. Natl. Acad. Sci. USA 109, 5568–5575 (2012).

17 Lang, A. H., Li, H., Collins, J. J. & Mehta, P. Epigenetic landscapes explain partially reprogrammed cells and identify key reprogramming genes. PLOS Comput. Biol. 10, e1003734 (2014).

18 Pusuluri, S. T., Lang, A. H., Mehta, P. & Castillo, H. E. Cellular reprogramming dynamics follow a simple 1D reaction coordinate. Phys. Biol. 15, 016001 (2017).

19 Weinreb, C., Wolock, S., Tusi, B. K., Socolovsky, M. & Klein, A. M. Fundamental limits on dynamic inference from single-cell snapshots. Proc. Natl. Acad. Sci. USA 115, E2467–E2476 (2018).

20 Jin, S., MacLean, A. L., Peng, T. & Nie, Q. scEpath: Energy landscape-based inference of transition probabilities and cellular trajectories from single-cell transcriptomic data. Bioinformatics 34, 2077–2086 (2018).

21 Marvel, S. A., Strogatz, S. H. & Kleinberg, J. M. Energy landscape of social balance. Phys. Rev. Lett. 103, 198701 (2009).

22 Facchetti, G., Iacono, G. & Altafini, C. Exploring the low-energy landscape of large-scale signed social networks. Phys. Rev. E 86, 036116 (2012).

23 Ballard, A. J. et al. Energy landscapes for machine learning. Phys. Chem. Chem. Phys. 19, 12585–12603 (2017).

24 Ferguson, A. L. et al. Translating HIV sequences into quantitative fitness landscapes predicts viral vulnerabilities for rational immunogen design. Immunity 38, 606–617 (2013).

25 Ebeling, W. & Feistel, R. Studies on Manfred Eigen’s model for the self-organization of information processing. Eur. Biophys. J. 47, 395–401 (2018).

26 Fischer, N., Konevega, A. L., Wintermeyer, W., Rodnina, M. V. & Stark, H. Ribosome dynamics and tRNA movement by time-resolved electron cryomicroscopy. Nature 466, 329–333 (2010).

27 Bai, X. C. et al. An atomic structure of human γ-secretase. Nature 525, 212–217 (2015).

28 Behrmann, E. et al. Structural snapshots of actively translating human ribosomes. Cell 161, 845–857 (2015).

29 Fernandez-Leiro, R. & Scheres, S. H. Unravelling biological macromolecules with cryo-electron microscopy. Nature 537, 339–346 (2016).

30 Frank, J. & Ourmazd, A. Continuous changes in structure mapped by manifold embedding of single-particle data in cryo-EM. Methods 100, 61–67 (2016).

31 Shalek, A. K. et al. Single-cell transcriptomics reveals bimodality in expression and splicing in immune cells. Nature 498, 236–240 (2013).

32 Kunegis, J., Lommatzsch, A. & Bauckhage, C. The Slashdot Zoo: Mining a social network with negative edges. Proceedings of the 18th International World Wide Web Conference (WWW’09), Madrid 741–750 (2009).

33 Leskovec, J., Huttenlocher, D. & Kleinberg, J. Signed networks in social media. Proc 28th CHI 1361 (2010).

34 Stephens, G. J., Osborne, L. C. & Bialek, W. Searching for simplicity in the analysis of neurons and behavior. Proc. Natl. Acad. Sci. USA 108, 15565–15571 (2011).

35 Chiavazzo, E. et al. Intrinsic map dynamics exploration for uncharted effective free-energy landscapes. Proc. Natl. Acad. Sci. USA 114, E5494–E5503 (2017).

36 Wasserman, L. Topological data analysis. Annu. Rev. Stat. Appl. 5, 501–532 (2018).

37 Mattingly, H. H., Transtrum, M. K., Abbott, M. C. & Machta, B. B. Maximizing the information learned from finite data selects a simple model. Proc. Natl. Acad. Sci. USA 115, 1760–1765 (2018).

38 Westerlund, A. M., Harpole, T. J., Blau, C. & Delemotte, L. Inference of Calmodulin’s Ca^2+^-dependent free energy landscapes via Gaussian mixture model validation. J. Chem. Theor. Comput. 14, 63–71 (2018).

39 Jónsson, H., Mills, G. & Jacobsen, K. W. Nudged elastic band method for finding minimum energy paths of transitions. In Classical and Quantum Dynamics in Condensed Phase Simulations, 385–404 (World Scientific, 1998).

40 Trygubenko, S. A. & Wales, D. J. A doubly nudged elastic band method for finding transition states. J. Chem. Phys. 120, 2082–2094 (2004).

41 Kusumaatmaja, H. Surveying the free energy landscapes of continuum models: Application to soft matter systems. J. Chem. Phys. 142, 124112 (2015).

42 Kramers, H. A. Brownian motion in a field of force and the diffusion model of chemical reactions. Physica 7, 284–304 (1940).

43 Malakhov, A. N. & Pankratov, A. L. Evolution times of probability distributions and averages - exact solutions of the Kramers’ problem. Adv. Chem. Phys. 121, 357–438 (2002).

44 Dunkel, J., Ebeling, W., Schimansky-Geier, L. & Hänggi, P. Kramers problem in evolutionary strategies. Phys. Rev. E 67, 061118 (2003).

45 van Kampen, N. G. Stochastic Processes in Physics and Chemistry (North-Holland Personal Library, Amsterdam, 2003).

46 Bowman, G. R. & Pande, V. S. Protein folded states are kinetic hubs. Proc. Natl. Acad. Sci. USA 107, 10890–10895 (2010).

47 Chodera, J. D. & Noé, F. Markov state models of biomolecular conformational dynamics. Curr. Opin. Struct. Biol. 25, 135–144 (2014).

48 Mardt, A., Pasquali, L., Wu, H. & Noé, F. VAMPnets for deep learning of molecular kinetics. Nat. Commun. 9, 5 (2018).

49 Lindorff-Larsen, K., Piana, S., Dror, R. O. & Shaw, D. E. How fast-folding proteins fold. Science 334, 517–520 (2011).

50 Sborgi, L. et al. Interaction networks in protein folding via atomic-resolution experiments and long-time-scale molecular dynamics simulations. J. Am. Chem. Soc. 137, 6506–6516 (2015).

51 Thomas, P., Popović, N. & Grima, R. Phenotypic switching in gene regulatory networks. Proc. Natl. Acad. Sci. USA 111, 6994–6999 (2014).

52 Schultz, D., Walczak, A. M., Onuchic, J. N. & Wolynes, P. G. Extinction and resurrection in gene networks. Proc. Natl. Acad. Sci. USA 105, 19165–19170 (2008).

53 Chu, B. K., Margaret, J. T., Sato, R. R. & Read, E. L. Markov State Models of gene regulatory networks. BMC Syst. Biol. 11, 14 (2017).

54 Kalai, A. T., Moitra, A. & Valiant, G. Disentangling Gaussians. Commun. ACM 55, 113–120 (2012).

55 Bühlmann, P., Kalisch, M. & Meier, L. High-dimensional statistics with a view toward applications in biology. Annu. Rev. Stat. Appl. 1, 255–278 (2014).

56 Lee, A. A., Brenner, M. P. & Colwell, L. J. Optimal design of experiments by combining coarse and fine measurements. Phys. Rev. Lett. 119, 208101 (2017).

57 Bolhuis, P. G. and Csányi, G. Nested transition path sampling. Phys. Rev. Lett. 120, 250601 (2018).

58 Müller, S. et al. A generalized model of the repressilator. J. Math. Biol. 53, 905–937 (2006).

59 Potvin-Trottier, L., Lord, N. D., Vinnicombe, G. & Paulsson, J. Synchronous long-term oscillations in a synthetic gene circuit. Nature 538, 514–517 (2016).

60 Ferrell Jr, J. E. Bistability, bifurcations, and Waddington’s epigenetic landscape. Curr. Biol. 22, R458–R466 (2012).

61 Zanini, F. et al. Population genomics of intrapatient HIV-1 evolution. eLife 4, e11282 (2015).

62 Gokhale, C. S., Iwasa, Y., Nowak, M. A. & Traulsen, A. The pace of evolution across fitness valleys. J. Theor. Biol. 259, 613–620 (2009).

63 Socci, N. D., Onuchic, J. N. & Wolynes, P. G. Diffusive dynamics of the reaction coordinate for protein folding funnels. J. Chem. Phys. 104, 5860–5868 (1996).

64 Zheng, W. & Best, R. B. Reduction of all-atom protein folding dynamics to one-dimensional diffusion. J. Phys. Chem. B. 119, 15247–15255 (2015).

65 Ceriotti, M., Tribello, G. A. & Parrinello, M. Simplifying the representation of complex free-energy landscapes using sketch-map. Proc. Natl. Acad. Sci. USA 108, 13023–13028 (2011).

66 Ferguson, A. L., Panagiotopoulos, A. Z., Kevrekidis, I. G. & Debenedetti, P. G. Nonlinear dimensionality reduction in molecular simulation: The diffusion map approach. Chem. Phys. Lett. 509, 1–11 (2011).

67 Das, P., Moll, M., Stamati, H., Kavraki, L. E. & Clementi, C. Low-dimensional, free-energy landscapes of protein-folding reactions by nonlinear dimensionality reduction. Proc. Natl. Acad. Sci. USA 103, 9885–9890 (2006).

68 Hegger, R., Altis, A., Nguyen, P. H. & Stock, G. How complex is the dynamics of peptide folding? Phys. Rev. Lett. 98, 028102 (2007).

69 Zhuravlev, P. I., Materese, C. K., Papoian, G. A. & Carolina, N. Deconstructing the native state: Energy landscapes, function, and dynamics of globular proteins. J. Phys. Chem. B. 113, 8800–8812 (2009).

70 Rohrdanz, M. A., Zheng, W. & Clementi, C. Discovering mountain passes via torchlight: methods for the definition of reaction coordinates and pathways in complex macromolecular reactions. Annu. Rev. Phys. Chem. 64, 295–316 (2013).

71 Krivov, S. V. On reaction coordinate optimality. J. Chem. Theor. Comput. 9, 135–146 (2013).

72 Best, R. B., Hummer, G. & Eaton, W. A. Native contacts determine protein folding mechanisms in atomistic simulations. Proc. Natl. Acad. Sci. USA 110, 17874–17879 (2013).

73 Ernst, M., Sittel, F. & Stock, G. Contact-and distance-based principal component analysis of protein dynamics. J. Chem. Phys..143, 244114 (2016).

74 Pressé, S., Ghosh, K., Lee, J. & Dill, K. A. Principles of maximum entropy and maximum caliber in statistical physics. Rev. Mod. Phys. 85, 1115 (2013).

75 Dixit, P. D., Jain, A., Stock, G. & Dill, K. A. Inferring transition rates of networks from populations in continuous-time Markov processes. J. Chem. Theor. Comput. 11, 5464–5472 (2015).

76 Dixit, P. D. et al. Perspective: Maximum caliber is a general variational principle for dynamical systems. J. Chem. Phys. 148, 010901 (2018).

77 Bar-Even, A. et al. Noise in protein expression scales with natural protein abundance. Nat. Genet. 38, 636–643 (2006).

78 Orr, H. A. Fitness and its role in evolutionary genetics. Nat. Rev. Genet. 10, 531–539 (2009).

79 Sella, G. & Hirsh, A. E. The application of statistical physics to evolutionary biology. Proc. Natl. Acad. Sci. USA 102, 9541–9546 (2005).

80 Barrick, J. E. & Lenski, R. E. Genome dynamics during experimental evolution. Nat. Rev. Genet. 14, 827–839 (2013).

81 Sharma, A., Wittmann, R. & Brader, J. M. Escape rate of active particles in the effective equilibrium approach. Phys. Rev. E 95, 012115 (2017).

