## Supplementary Information for "Learning dynamical information from static protein and sequencing data"

[1]*Department of Mathematics, Massachusetts Institute of Technology, Cambridge MA 02139-4307*
[2]*Mathematical Institute, University of Oxford, Andrew Wiles Building, Woodstock Road, Oxford OX2 6GG, United Kingdom*
[3]*Department of Physics, Durham University, Science Laboratories, South Road, Durham DH1 3LE, United Kingdom*
(Dated: May 21, 2019)

### I. POPULATION AND ENERGY LANDSCAPES

#### A. Gaussian mixture models

A Gaussian mixture model (GMM) was used to represent the probability density function (PDF), or population landscape, of samples. The PDF of a GMM with $C$ mixture components in $d$ dimensions is

$$p(\mathbf{x}) = \sum_{i=1}^{C} \phi_i p_i(\mathbf{x}), \tag{1}$$

where $\phi_i$ are the weights of each component and $\mathbf{x}$ is a $d$-dimensional vector. Each component in the mixture has a multivariate normal distribution with the $d$-dimensional mean vector $\boldsymbol{\mu}_i$ and the $d \times d$ covariance matrix $\boldsymbol{\Sigma}_i$. Therefore the PDF of each component is

$$p_i(\mathbf{x}) = \frac{\exp\left(-\frac{1}{2}\left(\mathbf{x}-\boldsymbol{\mu}_i\right)^T \boldsymbol{\Sigma}_i^{-1}\left(\mathbf{x}-\boldsymbol{\mu}_i\right)\right)}{\sqrt{\det\left(2\pi\boldsymbol{\Sigma}_i\right)}}. \tag{2}$$

The derivative of each multivariate normal in the mixture with respect to $\mathbf{x}$ is

$$\frac{\partial p_i(\mathbf{x})}{\partial \mathbf{x}} = -p_i(\mathbf{x}) \boldsymbol{\Sigma}_i^{-1}\left(\mathbf{x}-\boldsymbol{\mu}_i\right).$$

The Hessian of each multivariate normal in the mixture is

$$\frac{\partial^2 p_i(\mathbf{x})}{\partial \mathbf{x} \partial \mathbf{x}^T} = p_i(\mathbf{x})\left(\boldsymbol{\Sigma}_i^{-1}\left(\mathbf{x}-\boldsymbol{\mu}_i\right)\left(\mathbf{x}-\boldsymbol{\mu}_i\right)^T \boldsymbol{\Sigma}_i^{-1} - \boldsymbol{\Sigma}_i^{-1}\right).$$

The overall mean $\boldsymbol{\mu}$ and covariance $\boldsymbol{\Sigma}$ of a GMM are given by

$$\boldsymbol{\mu} = \sum_{i=1}^{C} \phi_i \boldsymbol{\mu}_i, \qquad \boldsymbol{\Sigma} = \sum_{i=1}^{C} \phi_i\left(\boldsymbol{\Sigma}_i + \boldsymbol{\mu}_i\left(\boldsymbol{\mu}_i - \boldsymbol{\mu}\right)\right). \tag{3}$$

Two Gaussians $N\left(\boldsymbol{\mu}_1;\ \sigma^2 \boldsymbol{I}_d\right)$ and $N\left(\boldsymbol{\mu}_2;\ \sigma^2 \boldsymbol{I}_d\right)$ in a mixture are *c-separated* if

$$|\boldsymbol{\mu}_1 - \boldsymbol{\mu}_2| \geq c\sigma\sqrt{d},$$

and a mixture of Gaussians is *c-separated* if the Gaussians in it are *c-separated*[1].

#### B. Artificial population landscapes

To construct artificial population landscapes for the mock data (Fig. 1), a Gaussian mixture model with 5 components in 10 dimensions was generated. The means were chosen to be 5 arbitrary points and the weights were chosen arbitrarily. The covariance matrices were arbitrary matrices satisfying the conditions necessary for our dimensionality reduction method, as described in the section "Preserving the topology of energy landscapes after dimensionality reduction" of the Supplementary Information.

#### C. Protein folding population landscapes

To construct population landscapes of proteins, we used the following algorithm. First, the number of Gaussians in the mixture was determined by calculating the optimal number of k-means clusters in the first two principal components of the data using the Calinski-Harabasz criterion, then adding four extra Gaussian components to increase the fit. This was found empirically to lead to accurate transition networks. Second, after the number of clusters had been determined, the clustering itself was performed by fitting a GMM in three dimensions using the expectation maximization (EM) algorithm; to reduce the number of fitted parameters, covariance matrices were assumed to be diagonal. Third, using the clusters that had been identified in three dimensions, a single Gaussian component was fit to each cluster of samples in the higher number of dimensions, again with diagonal covariance matrices; weights of each component in the mixture were given by the proportion of the total samples in the corresponding cluster. Population landscapes in five dimensions were found to be sufficient to capture the mean first passage times (Fig. S1A). The robustness of the results to different numbers of Gaussians in the mixture was tested by adding up to five extra Gaussians and recalculating transition networks between states identified on the original landscape (Fig. S1B). The error bars on Fig. S1B show the standard error calculated from these results.

#### D. Identification of protein states

The global minimum was designated to be the folded state and the second lowest minimum was designated to be the unfolded state. These classifications were confirmed to be accurate by calculating the root mean squared distance (RMSD) from the folded state obtained from the Protein Data Bank (https://www.rcsb.org/). An exception was WW, which had two minima with low RMSD; in this case, the folded and unfolded states were identified manually. Other low energy states were identified as minima on the energy landscape whose energy difference from the unfolded state was lower than the energy difference between the unfolded state and the folded state.

#### E. Data pre-processing: HIV sequences

HIV sequence data was obtained from a previous study, in which whole-genome deep sequencing of HIV-1 populations was performed in 9 untreated patients, with 6-12 samples per patient taken longitudinally at intervals over a period of 5-8 years[2]. Data from two of the patients (P4 and P7) was removed because they were excluded from the analysis in the original paper. A multi-sequence alignment (MSA) was performed on identified sequences in the p17 section of the HIV genome. Sequences were binarized using the method of Ref. 3, by comparing with the HIV type B consensus sequence identified by the Los Alamos National Laboratory HIV database (http://www.hiv.lanl.gov). Residues with nucleotides matching those of the consensus sequence were set to 0, and the remaining residues were set to 1 to denote a mutation. Sequences from the final 5 timepoints for each patient were retained to avoid bias by founder sequences and weighting towards patients with more longitudinal samples taken; at each timepoint the number of reads of each unique sequence was normalized by the amount of total reads at that timepoint. The first ten principle components of the discretized data were taken and assumed to correspond to a continuous phenotype space.

#### F. HIV population landscapes

A Gaussian was fit to binarized samples in the first 10 principal components for each patient. The GMM representing the population landscape was constructed by combining the individual Gaussians with equal weights.

#### G. GRN population landscapes

A Gaussian mixture model was fit to the $D$-dimensional protein-number data for each gene regulatory network (GRN) motif ($D = 2,\ 3,\ 4$). Owing to the low number of dimensions of this data, the number of minima was observable by inspection, and the number of Gaussians in the mixture was chosen to ensure each visible minimum was captured by the probably density function, as tested by the minima-finding algorithm.

### II. FINDING MINIMA AND MINIMUM ENERGY PATHWAYS

The energy landscape exploration involved locating the local minima in the system and the subseqent minimum energy pathways between them. To find the local minima we apply a random hopping algorithm. In this algorithm, a step consists of a trial move followed by an energy minimization. The simplest trial move consists of random perturbations for the system configuration. All unique local minima were accepted and stored.

The minimum energy pathway (MEP) between each pair of minima, through the corresponding saddle point, was calculated using the nudged elastic band (NEB) method[4]. The NEB method starts with the coordinates of two minima and attempts to trace out the MEP using a set of $N$ images with a set of cordinates $\{\boldsymbol{q_1}, \boldsymbol{q_2}, ..., \boldsymbol{q_N}\}$. A suitable initial guess is taken to be a linear path between the two minima, where the images at each end are fixed in place. Random perturbations to this initial guess were confirmed to give the same MEP. An image, with coordinates $\boldsymbol{q_n}$, is assumed to be connected to the two adjacent images, $\boldsymbol{q_{n-1}}$ and $\boldsymbol{q_{n+1}}$, via springs. The energy of the spring and image system is then minimised using a gradient calculated from the true gradient of the energy surface and the forces due to springs between the images, projected perpendicular and parallel to the vector tangent to the path, respectively, giving $\boldsymbol{g_n} = \boldsymbol{g}^{\text{true}}_{\boldsymbol{n},\perp} + \boldsymbol{g}^{\text{spring}}_{\boldsymbol{n},\parallel}$. Using the true and spring gradients without any projection is known to lead to corner-cutting (images are pulled away from the minimum energy path) and sliding-down problems (images slide down from barrier regions)[5].

In NEB, the relevant component of the gradient on an image at $\boldsymbol{q_n}$ due to the springs attached to adjacent images, $\boldsymbol{q_{n-1}}$ and $\boldsymbol{q_{n+1}}$ is the component parallel to the tangent vector $\boldsymbol{\tau_n}$. The projected spring gradient at image $n$, $\boldsymbol{g}^{\text{spring}}_{\boldsymbol{n}}$, is thus given by

$$\boldsymbol{g}^{\text{spring}}_{\boldsymbol{n},\parallel} = k\,\left(|\boldsymbol{q_{n+1}} - \boldsymbol{q_n}| - |\boldsymbol{q_{n-1}} - \boldsymbol{q_n}|\right)\hat{\boldsymbol{\tau}}_{\boldsymbol{n}} \tag{4}$$

with $k$ a tunable spring constant and the unit tangent vector $\hat{\boldsymbol{\tau}}_{\boldsymbol{n}} = \boldsymbol{\tau_n}/|\boldsymbol{\tau_n}|$. The tangent vector for image $n$ is calculated using the approach given in Ref. 5. If the image is neither at a maximum or a minimum then the tangent vector is defined as

$$\boldsymbol{\tau_n} = \begin{cases} \boldsymbol{\tau_{n,+}}, & \text{if } E_{n+1} > E_n > E_{n-1} \\ \boldsymbol{\tau_{n,-}}, & \text{if } E_{n+1} < E_n < E_{n-1} \end{cases} \tag{5}$$

where $E_n$ is the energy of image $n$ and $\boldsymbol{\tau_{n,\pm}} = \boldsymbol{q_{n\pm1}} - \boldsymbol{q_n}$. However, if the image $n$ is at a minimum or a maximum then the tangent is caculated using

$$\boldsymbol{\tau_n} = \begin{cases} \Delta E_{max}\boldsymbol{\tau_{n,+}} + \Delta E_{min}\boldsymbol{\tau_{n,-}}\ , & \text{if } E_{n+1} > E_{n-1} \\ \Delta E_{min}\boldsymbol{\tau_{n,+}} + \Delta E_{max}\boldsymbol{\tau_{n,-}}\ , & \text{if } E_{n+1} < E_{n-1} \end{cases} \tag{6}$$

where

$$\begin{aligned} \Delta E_{max} &= \max(|E_{n+1} - E_n|, |E_{n-1} - E_n|) \\ \Delta E_{min} &= \min(|E_{n+1} - E_n|, |E_{n-1} - E_n|). \end{aligned} \tag{7}$$

For the true gradient, in NEB we retain the component which is perpendicular to the unit tangent vector

$$\boldsymbol{g}^{\text{true}}_{\boldsymbol{n},\perp} = \boldsymbol{g}^{\text{true}}_{\boldsymbol{n}} - (\boldsymbol{g}^{\text{true}}_{\boldsymbol{n}} \cdot \hat{\boldsymbol{\tau}}_{\boldsymbol{n}})\hat{\boldsymbol{\tau}}_{\boldsymbol{n}}. \tag{8}$$

In some cases, owing to the finite spacing between the images, the local maxima found by NEB along the MEP deviate too far from the saddle points, so that our calculations for the Hessian and normal mode frequencies are inaccurate. To avoid this, we refine the saddle point position and energy by allowing the maximum energy image along the MEP to climb. We use the climbing image nudged elastic band method[6], which has been found to be highly succesful in accurately calculating saddle energies.

### III. KRAMERS RATES AND FIRST PASSAGE TIMES

#### A. Markov state models

Diffusion in a landscape comprising multiple minima, as in a Gaussian mixture model, can be coarse-grained into a discrete-state, continuous-time random walk between the basins of the minima. A direct transition between two minima is possible if they are connected by a minimum-energy pathway over a saddle point. If the minima are well separated and the energy barrier is sufficiently high, the transition waiting time is exponentially distributed with rate constant exponential in the energy barrier.

Formally, suppose we find $n$ minima with energies $E_1, \ldots, E_n$. The landscape algorithm finds all paths connecting pairs of minima via saddle points; let $S_{ij} = S_{ji}$ be the energy of the saddle point connecting minima $i$ and $j$, with $S_{ij} = \infty$ if there is no such saddle. The energy barrier for the direct transition $i \to j$ is then $\Delta E_{ij} = S_{ij} - E_i$. Under quadratic approximations for the basins and saddle, the waiting time for an overdamped diffusive transition $i \to j$ is distributed exponentially with Kramers rate constant[7]

$$k_{ij} = \frac{\omega_S}{2\pi\gamma} \frac{\prod_a \omega_a^i}{\prod_b' \omega_b^S} \exp\left(-\Delta E_{ij} / k_B T\right). \tag{9}$$

In this rate, $\omega_a^i$ are the $d$ angular frequencies of the energy $E$ at minimum $i$, while at the saddle $\omega_b^S$ are the $d-1$ stable angular frequencies and $\omega_S$ is the unstable angular frequency. This prefactor accounts for the shapes of minima and saddles, and is necessary to obtain the correct steady state distribution. Note that if $i$ and $j$ are not linked, with $S_{ij} = \infty$, then $k_{ij} = 0$.

Assuming transitions from one minimum to its neighbours are independent of one another, we can use the rates $k_{ij}$ to define a continuous-time Markov chain on the minima. Let $M_{ij} = k_{ij}$ be the transition rate from $i$ to $j$, and set $M_{ii} = -\sum_j k_{ij}$ so that rows sum to zero. The matrix $M$ is the generator matrix of such a chain.

Typically in diffusive systems a key quantity of interest is the mean first-passage time (MFPT) between a pair of states, such as folded and unfolded states of a protein. The MFPT $\tau_{ij}$, also called the hitting time, is the expected waiting time starting from state $i$ to first reach state $j$. For a fixed target $j$, the times for all start points $i$ can be computed by solving the linear system

$$\sum_i M_{ki}\tau_{ij} = -1 \quad \text{for } 1 \le k \le n,\ k \neq j,$$

$$\tau_{jj} = 0.$$

Solving the system for each $j$ then populates the entire pairwise MFPT matrix $\tau_{ij}$.

#### B. Quartic shape corrections

The Kramers rates in (9) assume that the neighbourhood geometries of the minima and saddles are well approximated as quadratics. For a GMM this is certainly true for the minima, but the saddles are 'pointy' with significant quartic terms in their expansion. As an example, consider the simplest symmetric case of a two-state one-dimensional symmetric GMM probability density $p(x) \propto e^{-(x-1)^2/2\sigma^2} + e^{-(x+1)^2/2\sigma^2}$. Then by symmetry the maximum of the energy $E(x)$ (which is the one-dimensional equivalent of a saddle point) between the minima $x \approx \pm 1$ is at $x = 0$, with expansion

$$\begin{aligned} E(x) &\equiv -\ln p(x) + \ln p_{\max} \\ &= \text{const} + \frac{\sigma^2 - 1}{2\sigma^4} x^2 + \frac{1}{12\sigma^8} x^4 + O(x^6). \end{aligned}$$

Assuming well separated minima with $\sigma \ll 1$, the quadratic term only dominates the quartic term on scales $x \ll \sigma^2 \lll 1$, suggesting a strong quartic influence on the rates. Indeed, the correction can be computed: from Ref. 7, the quartic saddle correction to (9) amounts to replacing $k_{ij} \to \phi k_{ij}$ where the prefactor $\phi$ reads

$$\phi = 1 - \frac{1}{8} \frac{E''''(0)}{E''(0)^2} = 1 - \frac{1}{4(\sigma^2 - 1)^2}.$$

Thus $\phi < 3/4$, so quartic corrections are numerically significant if calculating state transition rates from a full set of known parameters. That said, $\phi$ is approximately constant for $\sigma \ll 1$, approaching the limit $\phi \to 3/4$ as $\sigma \to 0$, so for a system of well-separated minima the contributions of quartic saddle corrections are likely to amount to a global correction factor which can be absorbed into a global effective friction constant.

#### C. Comparison with time-dependent transitions

Reference MFPTs were calculated using the time-dependent protein folding trajectories and GRN simulation data. The first passage time between state $i$ and state $j$ was defined as the time between entering state $i$ and entering state $j$, with a fixed lag time to reduce noise. The MFPT was calculated by taking the mean of the first passage times.

#### D. Comparison with maximum caliber method

An alternative set of MFPTs were calculated by supplying the size of the energy barriers to the maximum caliber method, which is described in detail elsewhere[8–10]. Specifically, using this method the transition rates $i \to j$ were taken to be[9]

$$k_{ij} = \mu \sqrt{\frac{\pi_j}{\pi_i}} \exp\left(-\gamma \Delta E_{ij}\right), \tag{10}$$

where $\pi_i$ is the stationary population of state $i$, $E_{ij}$ is the energy barrier between states $i$ and $j$, and $\mu$ and $\gamma$ are fitting parameters that we chose by minimizing the Euclidean distance between the reference and predicted vectors of MFPTs. Using these rates with the energy barriers calculated between states for the protein data, we found that the MFPTs predicted by maximum caliber did not capture both folding and unfolding transitions accurately, owing to the dependence of the MFPTs on the prefactors in Eq. (9) (Fig. S4).

### IV. GENE REGULATORY NETWORK SIMULATIONS

#### A. Method

We simulated three repressilator-type gene regulatory network motifs[11] with self-activation, in which each gene encodes a protein that activates the expression of its associated gene and represses another, with $D = 2$, 3 and 4 dimensions. The motifs were assumed to behave similarly to the mutual inhibition/self activation (MISA) system described in Ref. 12, which corresponds to the $D = 2$ case considered here.

Specifically, each gene (denoted A, B, C and D) encodes a transcription factor (protein), which forms homodimers[13] that can either bind to the promoter of another one of the genes, repressing its expression, or bind to the promoter of its own gene, activating its expression. Therefore, the promoter controlling each gene X can exist in either the unbound state $\mathrm{X}_{00}$, the activator-bound/repressor-unbound state $\mathrm{X}_{10}$, the activator-unbound/repressor-bound state $\mathrm{X}_{01}$, or the activator-bound/repressor-bound state $\mathrm{X}_{11}$. Each gene's associated protein x is produced at a rate $g_1$ in the activator-bound/repressor-unbound state and $g_0$ otherwise. Protein dimerization is assumed to occur simultaneously with binding to DNA.

All gene regulatory network motifs were simulated using a Gillespie stochastic simulation algorithm (SSA) in the SimBiology toolbox in Matlab. For brevity, below we list the reactions only for the $D = 4$ system; reactions and parameters for the $D = 2$ and $D = 3$ systems can be found in the simulation code available from Github (https://github.com/philip-pearce/learning-dynamical). Each simulation was initiated without transcription factors present, and with each promoter in the unbound state $\mathrm{X}_{00}$. In the $D = 4$ case, the parameters were taken to be $g_0 = 5\ \mathrm{s}^{-1}$, $g_1 = 14\ \mathrm{s}^{-1}$, $k = 1\ \mathrm{s}^{-1}$, $h_r = 10^{-4}\ \mathrm{s}^{-1}\mathrm{molecule}^{-2}$, $f_r = 10^{-2}\ \mathrm{s}^{-1}$, $h_a = 2\ \mathrm{s}^{-1}\mathrm{molecule}^{-2}$, $f_a = 10^{-1}\ \mathrm{s}^{-1}$ and the simulation was run for $10^7$ s. Deterministic ordinary differential equation (ODE) simulations of the same networks were performed using the ode15s solver in the SimBiology toolbox in Matlab with the same parameters and initial conditions as above. All reactions were assumed to obey the law of mass action[14].

#### B. Reactions

Protein synthesis

$$\mathrm{A}_{01} \xrightarrow{\mathrm{g}_0} \mathrm{A}_{01} + \mathrm{a}, \quad \mathrm{A}_{00} \xrightarrow{\mathrm{g}_0} \mathrm{A}_{00} + \mathrm{a}, \quad \mathrm{A}_{10} \xrightarrow{\mathrm{g}_1} \mathrm{A}_{10} + \mathrm{a}, \quad \mathrm{A}_{11} \xrightarrow{\mathrm{g}_0} \mathrm{A}_{11} + \mathrm{a}$$

$$\mathrm{B}_{01} \xrightarrow{\mathrm{g}_0} \mathrm{B}_{01} + \mathrm{b}, \quad \mathrm{B}_{00} \xrightarrow{\mathrm{g}_0} \mathrm{B}_{00} + \mathrm{b}, \quad \mathrm{B}_{10} \xrightarrow{\mathrm{g}_1} \mathrm{B}_{10} + \mathrm{b}, \quad \mathrm{B}_{11} \xrightarrow{\mathrm{g}_0} \mathrm{B}_{11} + \mathrm{b}$$

$$\mathrm{C}_{01} \xrightarrow{\mathrm{g}_0} \mathrm{C}_{01} + \mathrm{c}, \quad \mathrm{C}_{00} \xrightarrow{\mathrm{g}_0} \mathrm{C}_{00} + \mathrm{c}, \quad \mathrm{C}_{10} \xrightarrow{\mathrm{g}_1} \mathrm{C}_{10} + \mathrm{c}, \quad \mathrm{C}_{11} \xrightarrow{\mathrm{g}_0} \mathrm{C}_{11} + \mathrm{c}$$

$$\mathrm{D}_{01} \xrightarrow{\mathrm{g}_0} \mathrm{D}_{01} + \mathrm{d}, \quad \mathrm{D}_{00} \xrightarrow{\mathrm{g}_0} \mathrm{D}_{00} + \mathrm{d}, \quad \mathrm{D}_{10} \xrightarrow{\mathrm{g}_1} \mathrm{D}_{10} + \mathrm{d}, \quad \mathrm{D}_{11} \xrightarrow{\mathrm{g}_0} \mathrm{D}_{11} + \mathrm{d}$$

Protein degradation

$$\mathrm{a} \xrightarrow{\mathrm{k}} \phi, \quad \mathrm{b} \xrightarrow{\mathrm{k}} \phi, \quad \mathrm{c} \xrightarrow{\mathrm{k}} \phi, \quad \mathrm{d} \xrightarrow{\mathrm{k}} \phi$$

Repression

$$\mathrm{A_{00} + 2\,d \underset{f_r}{\overset{h_r}{\rightleftharpoons}} A_{01}}, \quad \mathrm{A_{10} + 2\,d \underset{f_r}{\overset{h_r}{\rightleftharpoons}} A_{11}},$$

$$\mathrm{B_{00} + 2\,a \underset{f_r}{\overset{h_r}{\rightleftharpoons}} B_{01}}, \quad \mathrm{B_{10} + 2\,a \underset{f_r}{\overset{h_r}{\rightleftharpoons}} B_{11}},$$

$$\mathrm{C_{00} + 2\,b \underset{f_r}{\overset{h_r}{\rightleftharpoons}} C_{01}}, \quad \mathrm{C_{10} + 2\,b \underset{f_r}{\overset{h_r}{\rightleftharpoons}} C_{11}},$$

$$\mathrm{D_{00} + 2\,c \underset{f_r}{\overset{h_r}{\rightleftharpoons}} D_{01}}, \quad \mathrm{D_{10} + 2\,c \underset{f_r}{\overset{h_r}{\rightleftharpoons}} D_{11}}$$

Activation

$$\mathrm{A_{00} + 2\,a \underset{f_a}{\overset{h_a}{\rightleftharpoons}} A_{10}}, \quad \mathrm{A_{01} + 2\,a \underset{f_a}{\overset{h_a}{\rightleftharpoons}} A_{11}},$$

$$\mathrm{B_{00} + 2\,b \underset{f_a}{\overset{h_a}{\rightleftharpoons}} B_{10}}, \quad \mathrm{B_{01} + 2\,b \underset{f_a}{\overset{h_a}{\rightleftharpoons}} B_{11}},$$

$$\mathrm{C_{00} + 2\,c \underset{f_a}{\overset{h_a}{\rightleftharpoons}} C_{10}}, \quad \mathrm{C_{01} + 2\,c \underset{f_a}{\overset{h_a}{\rightleftharpoons}} C_{11}},$$

$$\mathrm{D_{00} + 2\,d \underset{f_a}{\overset{h_a}{\rightleftharpoons}} D_{10}}, \quad \mathrm{D_{01} + 2\,d \underset{f_a}{\overset{h_a}{\rightleftharpoons}} D_{11}}$$

### V. BROWNIAN DYNAMICS SIMULATIONS

Brownian motion in a potential $U(\mathbf{x})$ was modeled by an overdamped Langevin equation[7]

$$\dot{\mathbf{x}} = -\frac{1}{\gamma}\nabla U(\mathbf{x}) + \sqrt{\frac{2k_B T}{\gamma}}\mathbf{R}(t), \tag{11}$$

where $\gamma$ is the friction, $k_B$ is Boltzmann's constant, $T$ is the temperature and $\mathbf{R}(t)$ is a delta-correlated stationary Gaussian process with zero-mean. (11) was simulated using a finite-difference approximation.

### VI. PRESERVING THE TOPOLOGY OF ENERGY LANDSCAPES AFTER DIMENSIONALITY REDUCTION

A Gaussian mixture model (GMM) in $D$ dimensions can be reduced to $d$ dimensions via the eigendecomposition of the covariance matrix of the Gaussian means

$$\boldsymbol{\Sigma} = \boldsymbol{U}\boldsymbol{D}\boldsymbol{U}^T. \tag{12}$$

Projection of a GMM defined by (1)-(3) onto the $d$-dimensional subspace spanned by the top $d$ eigenvectors of $\boldsymbol{\Sigma}$ gives a GMM with $C$ mixture components, which has the same mixing parameters $\phi_i$, means $\boldsymbol{\mu}_i^d = \boldsymbol{U}_d^T\boldsymbol{\mu}_i$ and variances $\boldsymbol{\Sigma}_i^d = \boldsymbol{U}_d^T\boldsymbol{\Sigma}_i\boldsymbol{U}_d$. Here $\boldsymbol{U}_d$ corresponds to the first $d$ columns of the matrix of sorted eigenvectors $\boldsymbol{U}$.

To reduce the dimension of an energy landscape while preserving its topology requires preserving the topology of the PDF in the reduced dimension. For each individual component of a GMM, the PDF at a point $\mathbf{x}$ in the original $D$-dimensional space can be calculated from the value of the PDF at a point $\mathbf{x}_d = \boldsymbol{U}_d^T\mathbf{x}$ in the reduced dimensional space as follows. In $d$ dimensions, inserting the projected coordinates, means and covariances into (2), the PDF is given by

$$p_i^d(\mathbf{x}_d) = \frac{\exp\left(-\frac{1}{2}\left[\boldsymbol{U}_d^T(\mathbf{x}-\boldsymbol{\mu}_i)\right]^T\left[\boldsymbol{U}_d^T\boldsymbol{\Sigma}_i\boldsymbol{U}_d\right]^{-1}\left[\boldsymbol{U}_d^T(\mathbf{x}-\boldsymbol{\mu}_i)\right]\right)}{\sqrt{\det 2\pi\boldsymbol{U}_d^T\boldsymbol{\Sigma}_i\boldsymbol{U}_d}}.$$

If we change coordinates to the basis of eigenvectors of $\boldsymbol{\Sigma}_i$ by substituting in $\boldsymbol{\Sigma}_i = \boldsymbol{S}\boldsymbol{D}\boldsymbol{S}^T$ with diagonal $\boldsymbol{D}$ and orthogonal $\boldsymbol{S}$, $\mathbf{y} = \boldsymbol{S}^T(\mathbf{x}-\boldsymbol{\mu}_i)$, and $\boldsymbol{V}_d = \boldsymbol{S}^T\boldsymbol{U}_d$, the terms in the exponent become

$$-\frac{1}{2}\left[\boldsymbol{U}_d^T(\mathbf{x}-\boldsymbol{\mu}_i)\right]^T\left[\boldsymbol{U}_d^T\boldsymbol{\Sigma}_i\boldsymbol{U}_d\right]^{-1}\left[\boldsymbol{U}_d^T(\mathbf{x}-\boldsymbol{\mu}_i)\right] = -\frac{1}{2}\mathbf{y}^T\boldsymbol{V}_d\left[\boldsymbol{V}_d^T\boldsymbol{D}\boldsymbol{V}_d\right]^{-1}\boldsymbol{V}_d^T\mathbf{y} \tag{13}$$

Suppose that $\boldsymbol{V}$ and $\mathbf{y}$ are zero outside of a subset of $d$ rows, i.e. that up to some permutation of the rows

$$\boldsymbol{V}_d = \begin{pmatrix} \boldsymbol{W} \\ \mathbf{0} \end{pmatrix} \qquad\qquad \mathbf{y} = \begin{pmatrix} \mathbf{y}^* \\ \mathbf{0} \end{pmatrix}$$

for some orthogonal matrix $\boldsymbol{W} \in \mathbb{R}^{d\times d}$ and $d$-dimensional vector $\mathbf{y}^*$. This is equivalent to requiring that the vector $(\mathbf{x} - \boldsymbol{\mu}_i)$ and the $d$ columns of $\boldsymbol{U}_d$ fall in a subspace spanned by $d$ eigenvectors of $\boldsymbol{\Sigma}_i$. Then (13) simplifies to

$$-\frac{1}{2}\mathbf{y}^T\boldsymbol{V}_d\left[\boldsymbol{V}_d^T\boldsymbol{D}\boldsymbol{V}_d\right]^{-1}\boldsymbol{V}_d^T\mathbf{y} = \mathbf{y}^{*T}\boldsymbol{D}^*\mathbf{y}^*,$$

where $\boldsymbol{D}^*$ is the submatrix of $\boldsymbol{D}$ restricted to the $d$ rows where $\boldsymbol{V}_d$ has nonzero entries. Note that $\boldsymbol{U}_d$ has completely dropped out of the equation. Under these circumstances, the contribution to the higher dimensional PDF from this component is related to the projected PDF by a simple scaling:

$$p_i^d\left(\mathbf{x}_d\right) = p_i(\mathbf{x})\frac{\sqrt{\det\left(2\pi\boldsymbol{\Sigma}_i\right)}}{\sqrt{\det\left(2\pi\boldsymbol{U}_d^T\boldsymbol{\Sigma}_i\boldsymbol{U}_d\right)}},$$

with no change to the exponent.

Therefore, if the value of the PDF is known at a point $\mathbf{x}_d$ in $d$-dimensional PC-space, the corresponding value of the PDF in the original $D$-dimensional space can be calculated using

$$p\left(\mathbf{x}\right) = \sum_{i=1}^{C}\phi_i p_i^d\left(\mathbf{x}_d\right)\frac{\sqrt{\det\left(2\pi\boldsymbol{U}_d^T\boldsymbol{\Sigma}_i\boldsymbol{U}_d\right)}}{\sqrt{\det\left(2\pi\boldsymbol{\Sigma}_i\right)}}. \tag{14}$$

For a mixture of spherical Gaussians with covariance matrices $\boldsymbol{\Sigma}_i = \sigma_i\mathbf{I}$, equation (14) becomes

$$p\left(\mathbf{x}\right) = \sum_{i=1}^{C}\phi_i p_i^d\left(\mathbf{x}_d\right)\left(\frac{1}{\sqrt{2\pi\sigma_i}}\right)^{D-d}, \tag{15}$$

and in this specific case a GMM needs only to be fit to the data in the reduced dimensional space.

(14) can be used to find MEPs in a low-dimensional space before converting back to the high-dimensional space and calculating Hessian matrices for substitution into (9) for the Kramers rates.

### VII. FINITE SAMPLE EFFECTS

In applications, we must estimate the energy landscapes using a finite number of samples from the underlying distribution. To understand how sampling error affects our calculations, we compute here the expectation and variance of the maximum likelihood estimate of the barrier and minima energies in a simplified case with two well-separated Gaussians with known, equal covariance matrices $\Sigma_1 = \Sigma_2$ and known, equal mixing weights $\phi_1 = \phi_2 = \frac{1}{2}$. The full PDF is

$$p(\mathbf{x}) = \phi_1\frac{1}{\sqrt{(2\pi)^d|\Sigma_1|}}e^{-\frac{1}{2}(\mathbf{x}-\mu_1)^\top\Sigma_1^{-1}(\mathbf{x}-\mu_1)} + \phi_2\frac{1}{\sqrt{(2\pi)^d|\Sigma_2|}}e^{-\frac{1}{2}(\mathbf{x}-\mu_2)^\top\Sigma_2^{-1}(\mathbf{x}-\mu_2)}. \tag{16}$$

If the Gaussians are well-separated, direct transitions between two minima will approximately follow the straight line between the two means. Here $\mathbf{x} = \mu_1 + c(\mu_2 - \mu_1)$, and we can reduce to a one-dimensional problem

$$p(\mathbf{x}(c)) = \phi_1\frac{1}{\sqrt{(2\pi)^d|\Sigma_1|}}e^{-\frac{s_1}{2}c^2} + \phi_2\frac{1}{\sqrt{(2\pi)^d|\Sigma_2|}}e^{-\frac{s_2}{2}(1-c)^2}. \tag{17}$$

where $s_1 = (\mu_2 - \mu_1)^\top\Sigma_1^{-1}(\mu_2 - \mu_1)$ and $s_2 = (\mu_1 - \mu_2)^\top\Sigma_2^{-1}(\mu_1 - \mu_2)$. The barrier is at the minimum as a function of $c$, where

$$0 = \partial_c p(c) = -\phi_1\frac{s_1 c}{\sqrt{(2\pi)^d|\Sigma_1|}}e^{-\frac{s_1}{2}c^2} + \phi_2\frac{s_2(1-c)}{\sqrt{(2\pi)^d|\Sigma_2|}}e^{-\frac{s_2}{2}(1-c)^2}. \tag{18}$$

As we assumed $\phi_1 = \phi_2 = \frac{1}{2}$ and $\Sigma_1 = \Sigma_2$, we have $s_1 = s_2 = s$ and the minimum occurs at $c = 1/2$, with a barrier probability

$$\frac{1}{\sqrt{(2\pi)^d|\Sigma_1|}}e^{-\frac{s}{8}}. \tag{19}$$

With real data, we need to deal with estimates from a finite sample. Suppose that we are able to cluster perfectly, i.e., that we know which Gaussian each sample came from. The only remaining parameters to estimate are the two means. The maximum likelihood estimates are the empirical means,

$$\hat{\mu}_1 = \frac{1}{n_1}\sum x_i^{(1)}, \qquad \hat{\mu}_2 = \frac{1}{n_2}\sum x_i^{(2)}$$

where $n_1$, $n_2$ are the sample sizes for groups 1 and 2 respectively, with corresponding samples $x_i^{(1)}$ and $x_i^{(2)}$. These estimators have empirical distribution

$$\hat{\mu}_1 \sim \mathcal{N}(\mu_1, \Sigma_1/n_1), \qquad \hat{\mu}_2 \sim \mathcal{N}(\mu_2, \Sigma_2/n_2)$$

Since we've assumed the variances are known, the true probabilities at the means $\mu_i$ will nearly match the fitted probabilities at the estimated means $\hat{\mu}_i$. The minima, for well separated Gaussians, will be approximately at the means, so we can estimate the minima energy well. For the barriers, with $n_1 = n_2 = n$, we get an estimate of the barrier probability

$$\hat{p}_b = \frac{1}{\sqrt{(2\pi)^d|\Sigma_1|}}e^{-\frac{\hat{s}}{8}},$$

where $\hat{s} = (\hat{\mu}_1 - \hat{\mu}_2)^\top \Sigma_1^{-1}(\hat{\mu}_1 - \hat{\mu}_2)$ is the estimated Mahalanobis distance between the means. This corresponds to an energy

$$\begin{aligned}
\hat{E}_b &= -\log(\hat{p}_b) \\
&= \log\left(\sqrt{(2\pi)^d|\Sigma_1|}\right) + \frac{\hat{s}}{8} \\
&= \log\left(\sqrt{(2\pi)^d|\Sigma_1|}\right) + \frac{(\hat{\mu}_1 - \hat{\mu}_2)^\top \Sigma_1^{-1}(\hat{\mu}_1 - \hat{\mu}_2)}{8} \\
&= \log\left(\sqrt{(2\pi)^d|\Sigma_1|}\right) + \frac{(\mu_1 - \mu_2 + \epsilon)^\top \Sigma_1^{-1}(\mu_1 - \mu_2 + \epsilon)}{8} \\
&= E_b + \frac{\epsilon^\top \Sigma_1^{-1}(\mu_1 - \mu_2)}{4} + \frac{\epsilon^\top \Sigma_1^{-1}\epsilon}{8}
\end{aligned}$$

where the error term $\epsilon = (\hat{\mu}_1 - \hat{\mu}_2) - (\mu_1 - \mu_2)$ has distribution $\mathcal{N}(0, 2\Sigma_1/n)$. This has expectation

$$\begin{aligned}
\mathbb{E}\left[\hat{E}_b\right] &= E_b + \frac{1}{8}\mathbb{E}\left[\epsilon^\top \Sigma_1^{-1}\epsilon\right] \\
&= E_b + \frac{d}{4n}
\end{aligned} \tag{20}$$

and variance

$$\begin{aligned}
\mathbb{E}\left[\left(\hat{E}_b - E_b - \frac{d}{4n}\right)^2\right] &= \frac{1}{64}\mathbb{E}\left[\left(2\epsilon^\top\Sigma_1^{-1}(\mu_1-\mu_2) + \epsilon^\top\Sigma_1^{-1}\epsilon - 2d/n\right)^2\right] \\
&= \frac{1}{64}\mathbb{E}\left[(2\epsilon^\top\Sigma_1^{-1}(\mu_1-\mu_2))^2\right. \\
&\qquad + 4(\epsilon^\top\Sigma_1^{-1}(\mu_1-\mu_2))(\epsilon^\top\Sigma_1^{-1}\epsilon - 2d/n) \\
&\qquad \left. +(\epsilon^\top\Sigma_1^{-1}\epsilon - 2d/n)^2\right] \\
&= \frac{1}{64}\mathbb{E}\left[(2\epsilon^\top\Sigma_1^{-1}(\mu_1-\mu_2))^2\right. \\
&\qquad \left. +(\epsilon^\top\Sigma_1^{-1}\epsilon - 2d/n)^2\right] \\
&= \frac{1}{64}\mathrm{tr}(4\Sigma_1^{-1}(\mu_1-\mu_2)(\mu_1-\mu_2)^\top\Sigma_1^{-1}\mathbb{E}[\epsilon\epsilon^\top]) \\
&\qquad + \frac{1}{64}\mathbb{E}\left[(\epsilon^\top\Sigma_1^{-1}\epsilon - 2d/n)^2\right] \\
&= \frac{1}{8}(\mu_1-\mu_2)^\top\Sigma_1^{-1}(\mu_1-\mu_2)/n \\
&\qquad + \frac{1}{64}\mathrm{Var}\left(\epsilon^\top\Sigma_1^{-1}\epsilon\right) \\
&= \frac{s}{8n} + \frac{1}{16n^2}\mathrm{Var}\left(\epsilon^\top(2\Sigma_1/n)^{-1}\epsilon\right).
\end{aligned}$$

In the above calculation, all the odd moments are zero since $\epsilon$ is Gaussian, and the quadratic moments can be calculated by taking traces, e.g.

$$\mathbb{E}[\epsilon^\top\Sigma_1^{-1}\epsilon] = \mathbb{E}[\mathrm{tr}(\Sigma_1^{-1}\epsilon\epsilon^\top)] = \mathrm{tr}(\Sigma_1^{-1}\mathbb{E}[\epsilon\epsilon^\top]) = \mathrm{tr}(\Sigma_1^{-1}(2\Sigma_1/n)) = 2d/n. \tag{21}$$

For the variance of the final term, we can decompose $\epsilon^\top(2\Sigma_1/n)^{-1}\epsilon$ into a sum of squares of independent normally-distributed variables using the eigenvalue decomposition of $\Sigma_1$; this is a $\chi^2$ distribution with $d$ degrees of freedom, with variance $2d$. So

$$\mathrm{Var}(\hat{E}_b) = \frac{s}{8n} + \frac{d}{8n^2} \tag{22}$$

In comparison, the true energy barrier height above the minima (approximated as the mean of one Gaussian) is

$$E_b - E_{\min} \approx \log\left(\sqrt{(2\pi)^d|\Sigma_1|}\right) + \frac{s}{8} - \log\left(\sqrt{(2\pi)^d|\Sigma_1|}\right) = \frac{s}{8}. \tag{23}$$

Unless $s$, the Mahalanobis distance between means, grows with dimension, in high dimensions with $d \gg n$ the variance will dominate and it will be impossible to accurately estimate the barrier height. If the coordinates of the means are independent and identically distributed, then the distance between means $s$ should scale with $\sqrt{d}$. So for a fixed sample size $n$, $\mathrm{Var}(\Delta\hat{E}) \sim \left(\mathbb{E}\left[\Delta\hat{E}\right]\right)^2$ and the energy barrier estimates will typically be off by roughly constant factors.

This provides an approximate lower bound on the number of samples required for recovering energy barriers. In addition, for moderate sample sizes $n \sim d$ there will be a bias in the barrier heights from (20) that must be corrected.

---


[1] Dasgupta, S. Learning mixtures of gaussians. In *Foundations of computer science, 1999. 40th annual symposium on*, 634–644 (IEEE, 1999).

[2] Zanini, F. *et al.* Population genomics of intrapatient HIV-1 evolution. *eLife* **4**, e11282 (2015).

[3] Ferguson, A. L. *et al.* Translating HIV Sequences into Quantitative Fitness Landscapes Predicts Viral Vulnerabilities for Rational Immunogen Design. *Immunity* **38**, 606–617 (2013).

[4] Jónsson, H., Mills, G. & Jacobsen, K. W. Nudged elastic band method for finding minimum energy paths of transitions. In *Classical and Quantum Dynamics in Condensed Phase Simulations*, 385–404 (World Scientific, 1998).

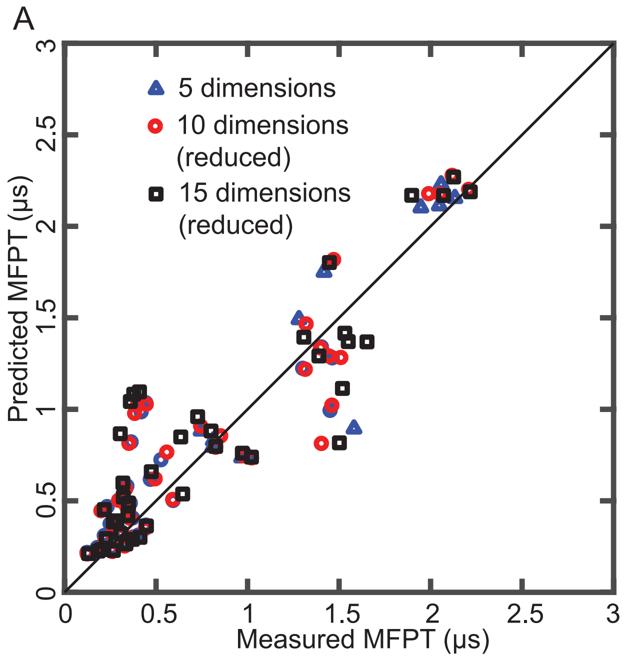


FIG. S1. *(A)* Comparison between predicted and measured MFPTs in 5D, 10D and 15D for villin, demonstrating that the first 5 principal components are enough to capture the transition network accurately. The 10D and 15D results were obtained after reducing to 7D using (14). *(B)* Comparison between predicted and measured MFPTs in 5D for villin, for different numbers of Gaussians in the mixture. The markers show the mean MFPT after adding up to five extra Gaussians and recalculating transition networks between states identified on the original landscape each time. Error bars show the standard error in the calculation.


[5] Henkelman, G. & Jónsson, H. Improved tangent estimate in the nudged elastic band method for finding minimum energy paths and saddle points. *J. Chem. Phys.* **113**, 9978–9985 (2000).

[6] Henkelman, G., Uberuaga, B. P. & Jónsson, H. A climbing image nudged elastic band method for finding saddle points and minimum energy paths. *J. Chem. Phys.* **113**, 9901–9904 (2000).

[7] Hänggi, P., Talkner, P. & Borkovec, M. Reaction-rate theory: Fifty years after Kramers. *Rev. Mod. Phys.* **62**, 251 (1990).

[8] Pressé, S., Ghosh, K., Lee, J. & Dill, K. A. Principles of maximum entropy and maximum caliber in statistical physics. *Rev. Mod. Phys.* **85**, 1115 (2013).

[9] Dixit, P. D., Jain, A., Stock, G. & Dill, K. A. Inferring transition rates of networks from populations in continuous-time Markov processes. *J. Chem. Theory Comput.* **11**, 5464–5472 (2015).

[10] Dixit, P. D. *et al.* Perspective: Maximum caliber is a general variational principle for dynamical systems. *J. Chem. Phys.* **148**, 010901 (2018).

[11] Müller, S. *et al.* A generalized model of the repressilator. *J. Math. Biol.* **53**, 905–937 (2006).

[12] Chu, B. K., Margaret, J. T., Sato, R. R. & Read, E. L. Markov State Models of gene regulatory networks. *BMC Syst. Biol.* **11**, 14 (2017).

[13] Funnell, A. P. & Crossley, M. Homo-and heterodimerization in transcriptional regulation. In *Protein Dimerization and Oligomerization in Biology*, 105–121 (Springer, 2012).

[14] Murray, J. D. *Mathematical Biology* (Springer-Verlag, 2003).

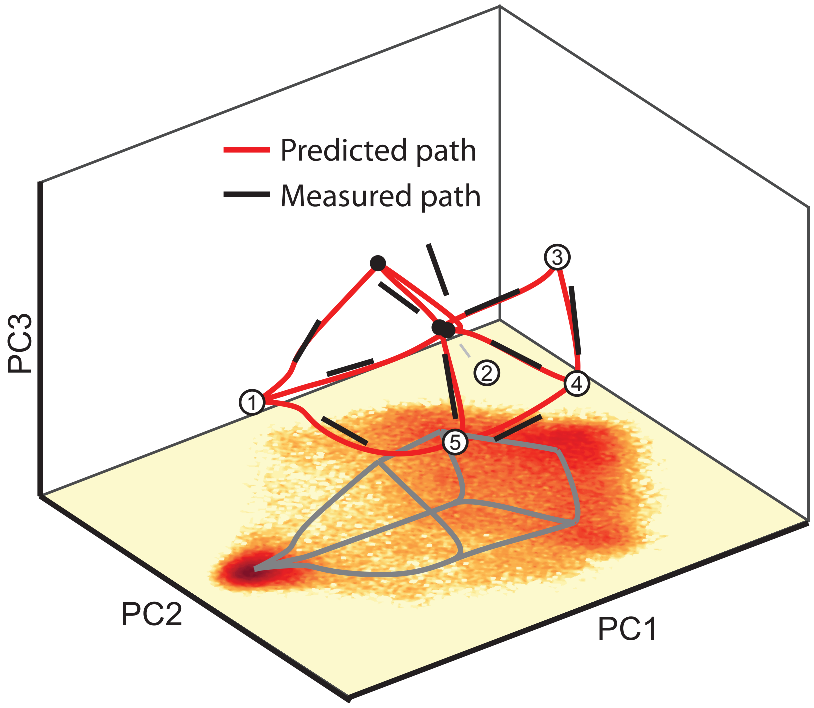


FIG. S2. States and transition network in the first three principal components (PCs) for villin including predicted transition paths between states (red lines) and transition paths from time-dependent data (black lines). Transition paths were calculated from time-dependent molecular dynamics (MD) data by drawing a straight line between the average pre-transition point and the average post-transition point, where a transition was defined as a change in cluster from one time-point to the next (see "Protein folding population landscapes" for a description of how clusters were identified). The two-dimensional projection of the empirical energy landscape onto the first two PCs is shown for illustration.

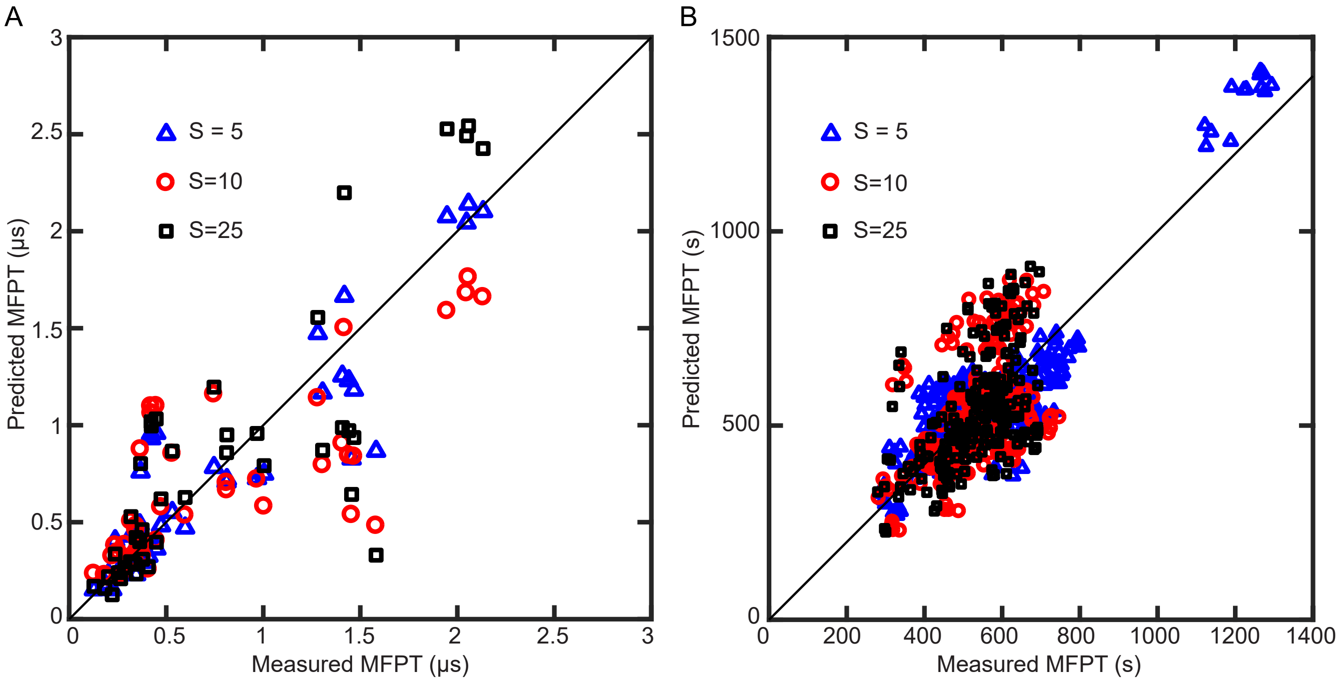


FIG. S3. *(A)* Comparison between predicted and measured MFPTs in 5D for villin after further subsampling the data by a factor $S = 5$, 10 or 25, leading to around 2500, 1300 or 500 samples per Gaussian, respectively. The dataset used in the main paper (see Fig. 2) was already subsampled from the available data by a factor of 5 and consisted of approximately $10^5$ samples. *(B)* Comparison between predicted and measured MFPTs in 4D for the gene regulatory network example after further subsampling the data by a factor $S = 5$, 10 or 25, leading to around $7 \cdot 10^3$, $3 \cdot 10^3$ or $1 \cdot 10^3$ samples per Gaussian, respectively. The energy landscapes constructed from the $S = 10$ and $S = 25$ datasets do not capture one of the highest energy minima (specifically, the minimum corresponding to low quantities of all four proteins), but the MFPTs are still accurate. The dataset used in the main paper (see Fig. 3) was already subsampled from the available data by a factor of $10^3$ and consisted of approximately $8 \cdot 10^5$ samples.

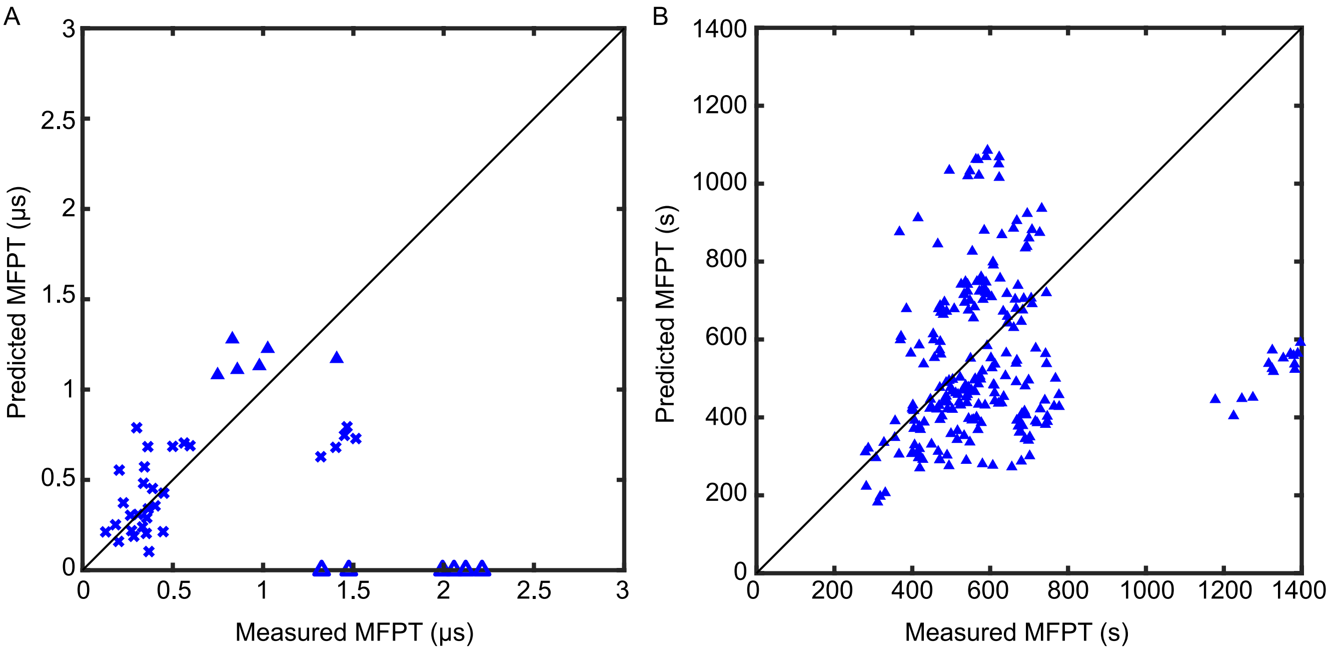


FIG. S4. *(A)* Comparison between predicted and measured MFPTs in 5D for villin, after supplying the sizes of the energy barriers to the maximum caliber method[8–10]. Filled triangles correspond to unfolding transitions and unfilled triangles correspond to folding transitions. The Pearson correlation coefficient between predicted and measured MFPTs is $\rho = -0.03$ for the maximum caliber method, in comparison to $\rho = 0.91$ for our method. *(B)* Comparison between predicted and measured MFPTs in 4D for the repressilator-type gene regulatory network shown in Fig. 3, after supplying the sizes of the energy barriers to the maximum caliber method. The Pearson correlation coefficient between predicted and measured MFPTs is $\rho = 0.40$ for the maximum caliber method, in comparison to $\rho = 0.94$ for our method. In both cases, because of the variation in the prefactors between transitions in Eq. (9), the maximum caliber method is unable to predict all transitions accurately.
